# Core Protein-Directed Antivirals and Importin β Can Synergistically Disrupt HBV Capsids

**DOI:** 10.1101/2021.08.16.456586

**Authors:** Christine Kim, Lauren Barnes, Christopher Schlicksup, Angela Patterson, Brian Bothner, Martin Jarrold, Che-Yen Joseph Wang, Adam Zlotnick

**Affiliations:** Department of Molecular & Cellular Biochemistry, Indiana University; Department of Chemistry, Indiana University; Department of Chemistry & Biochemistry, Montana State University; Department of Microbiology & Immunology, Pennsylvania State University College of Medicine

## Abstract

Viral structural proteins can have multiple activities. Antivirals that target structural proteins have potential to exhibit multiple antiviral mechanisms. Hepatitis B Virus (HBV) core protein (Cp) is involved in most stages of the viral lifecycle: it assembles into capsids, packages viral RNA, is a metabolic compartment for reverse transcription, interacts with nuclear trafficking machinery, and disassembles to release the viral genome into the nucleus. During nuclear localization, HBV capsids bind to host importins (e.g. Impβ) via Cp’s C-terminal domain (CTD); the CTD is localized to the interior of the capsid and is transiently exposed on the exterior. We used HAP12 as a representative Cp Allosteric Modulators (CpAMs), a class of antivirals that inappropriately stimulates and misdirects HBV assembly and deforms capsids. CpAM impact on other aspects of the HBV lifecycle is poorly understood. We investigated how HAP12 influenced the interactions between empty or RNA-filled capsids with Impβ and trypsin *in vitro*. We showed that HAP12 can modulate CTD accessibility and capsid stability, depending on the saturation of HAP12-binding sites. We demonstrated that Impβ synergistically contributes to capsid disruption at high levels of HAP12 saturation, using electron microscopy to visualize disruption and rearrangement of Cp dimers into aberrant complexes. However, RNA-filled capsids resisted the destabilizing effects of HAP12 and Impβ. In summary, we show host protein-induced catalysis of capsid disruption, an unexpected additional mechanism of action for CpAMs. Potentially, untimely capsid disassembly can hamper the HBV lifecycle and also cause the virus to become vulnerable to host innate immune responses.

**IMPORTANCE:** The HBV core, an icosahedral complex of 120 copies of the homodimeric core (capsid) protein with or without packaged nucleic acid, is transported to the host nucleus by its interaction with host importin proteins. Importin-core interaction requires the core protein C-terminal domain, which is inside the capsid, to “flip” to the capsid exterior. Core-protein directed drugs that affect capsid assembly and stability have been developed recently. We show that these molecules can, synergistically with importins, disrupt capsids. This mechanism of action, synergism with host protein, has potential to disrupt the virus lifecycle and activate the innate immune system.

## INTRODUCTION

Chronic Hepatitis B Virus (HBV) infection is endemic and, though not a regular discussion in the daily news, a global health crisis (1). Chronic HBV afflicts approximately 300 million people and can lead to cirrhosis, hepatocellular carcinoma, and liver failure; it contributes to about 880,000 deaths annually (2). Although there is an effective vaccine available, it does not help those who are chronically infected. Current therapeutics (mainly directed against the viral DNA polymerase) are rarely curative, so there is a great need to develop new and better antivirals.

An attractive drug target for new HBV therapeutics is the core protein (Cp) (3). The Cp plays roles in most stages of the viral lifecycle: assembling on and encapsidating viral RNA and polymerase, acting as a metabolic compartment for reverse transcription to DNA, and regulating capsid transport to the nucleus to maintain infection or to the ER to be secreted (4–6). Most capsids are composed of 120 homodimers arranged with T=4 icosahedral symmetry. A small fraction of capsids have 90 dimers with T=3 symmetry. Dimers associate through weak hydrophobic contacts at the inter-dimer interfaces (7). Cp has an assembly domain (residues 1-149) and a nucleic acid-binding C-terminal domain (CTD, residues 150-183). During viral replication, approximately 90% of capsids are without a viral genome (8). In T=4 capsids, Cp is found as quasi-equivalent A, B, C, and D monomers, which form AB and CD dimers. The interface between two dimers forms a small hydrophobic pocket, which can be probed with small molecules (4, 9).

Core protein allosteric modulators (CpAMs) are small molecules that can probe HBV capsids by binding to interdimer contacts (10). Heteroaryldihydropyrimidines (HAPs) are a class of CpAMs that have been extensively studied for their ability to accelerate and misdirect capsid assembly (11–13). When bound at Cp-Cp contact sites, in the HAP pocket, HAPs increase the association energies of dimer-dimer contacts. Cp assembled in the presence of HAPs can produce aberrant structures with varying morphology, depending on the CpAM chemotype (13–15). HAPs can also cause capsid deformation by disturbing the capsid’s icosahedral geometry. When incubated with the molecule HAP-TAMRA, capsid quasi-sixfold vertices became flattened, a defect that can propagate to yield highly irregular particles (15). Thus, not only do these antivirals impair capsid assembly, but they can also target morphologically “normal” capsids, even “melting” virions to prevent infection (16, 17). Although we understand how HAPs influence the capsid’s structure, it is still poorly understood how HAP-induced deformation impacts HBV biology in an infected cell.

Much of HBV Cp biology is a function of the CTDs. The CTDs are intrinsically disordered and positively charged (16 arginines out of 34 residues). In the context of an icosahedral capsid, CTDs are clustered around quasi-sixfold and five-fold vertices (18). CTDs play important roles in RNA packaging and regulating reverse transcription (19–21). Though they are ostensibly on the inside of the capsid, they can transiently flip out to the capsid exterior to expose nuclear localization signals (22–26). The organization and mobility of CTDs are influenced by phosphorylation and nucleic acid content (27–30). In the nucleic acid-filled particles, CTDs will electrostatically interact with the negatively charged nucleic acid and stay primarily internalized (18). During infection, capsids, containing relaxed circular DNA (rcDNA), can localize to the nucleus via interaction with host importins α (Impα and β (Impβ) (31). Once the capsid is imported into the nuclear pore complex, the capsid protein will interact with nucleoporin 153, disassemble, and release its genome (32, 33). Previously, it was shown that both Impα and Impβ were required for nuclear trafficking (33–35). However, Impβ can directly bind to the empty capsids or dimers without the presence of Impα and can be internalized (26).

Here, we examine how CpAMs affect CTD mobility. How these antivirals impact other aspects of the lifecycle is still unclear. For these studies we use the CpAM HAP12, which has strong Cp-binding activity in vitro and efficacy in vivo (13). HAP12 is structurally very similar to GLS4 (12). In this study, we investigated how HAP12-binding affects the CTD’s ability to interact with Impβ. We found that the CTD’s mobility increased with high HAP12 stoichiometry, causing more CTD-Impβ-binding. Surprisingly, excess HAPs led to capsid disruption in empty particles. However, despite excess HAPs, capsids with internal RNA did not experience greater Impβ-binding and were able to maintain their structural integrity. In this work, we showed how two exogenous molecules work synergistically to destabilize HBV capsids. Our study also investigated how a second antiviral mechanism of action, capsid dissociation, is modulated by CpAMs and host proteins.

## RESULTS

### Capsid proteolysis and Impβ binding experimental schematic

HAPs, like many CpAMs, cause capsid deformation and even disruption on a global level (14, 15). Because Cp dimers appear to be relatively rigid and CpAMs bind at the interface between dimers, deformation is likely to be manifested in the dimer-dimer geometry and at vertices. CTDs are clustered around fivefold and quasi-sixfolds vertices, therefore, we anticipated that HAPs would change the external exposure of CTDs.

To test our prediction that CpAMs modulate CTD exposure, we determined the effect of the CpAM HAP12 on CTD susceptibility to proteolysis and CTD ability to bind Impβ (Figure 1A and 1B). Impβ binds a basic peptide of ca. 40 amino acids (36), much longer than a typical nuclear localization sequence (NLS); therefore, Impβ is a sensitive tool for probing CTD exposure. The impact of CpAMs on CTD exposure has not previously been tested. Complicating this test, the effect of a HAP on assembly is highly dose dependent. At low ratios of HAP to Cp, morphologically normal capsids form while when HAP is super-stoichiometric – that is, there is more than one active HAP per subunit – abnormal polymers assemble (13, 14). For this reason, we wanted to examine how different ratios of CpAM modulated CTD exposure.

**Figure 1.**
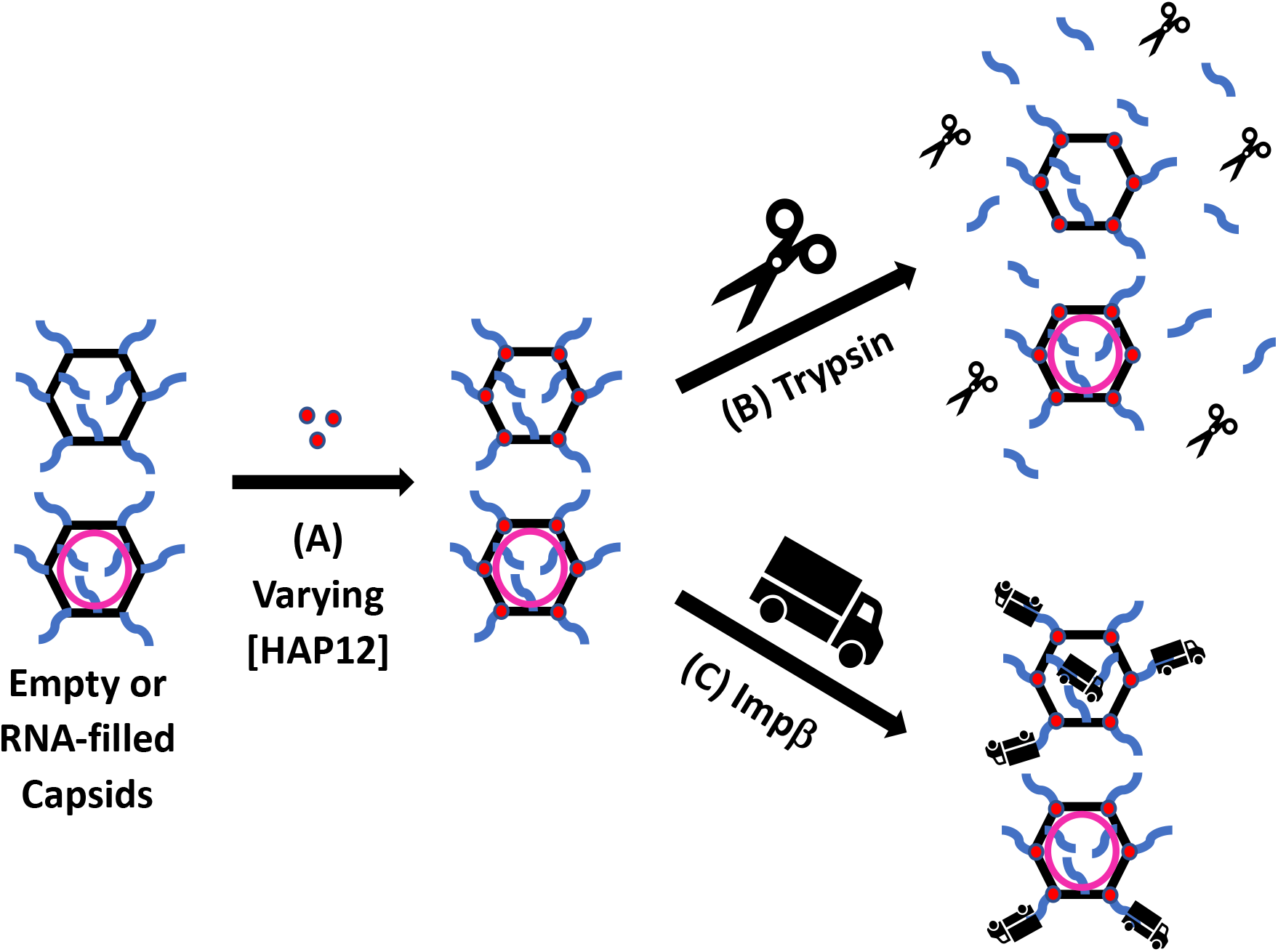
Capsid proteolysis and Impβ binding experimental schematic. (A) Empty and RNA-filled capsids were initially treated with varying concentrations of the CpAM HAP12 and then interrogated by proteolysis or Impβ-binding. Trypsin is represented by scissors, and Impβ is represented by trucks. HAP12-treatment was characterized in terms of “saturation”: at saturating HAP12 concentrations, there is one active HAP12 molecule for every HAP pocket. (B) HAP12-treated empty, pgRNA-filled, and E. coli RNA-filled capsids were digested with trypsin and reactions were quenched before proteolyzed products were resolved using SDS-PAGE and LC-MS. (C). HAP12-treated empty and E. coli RNA-filled capsids were mixed with Impβ and dialyzed into a low NaCl concentration buffer to allow binding. Impβ–bound capsids were analyzed using SEC, SDS-PAGE, TEM, and CDMS.

### High concentrations of the CpAM HAP12 change the CTD proteolysis pattern

Low HAP12:dimer ratios stabilize empty capsids and were predicted to decrease CTD proteolysis. Conversely, super-saturating ratios of HAP12, which cause capsid deformation even with intact capsids, were predicted to make empty capsids more vulnerable to proteolysis. To test this, empty capsids were treated with HAP12, proteolyzed with trypsin, and the digested products were analyzed by SDS-PAGE and LC-MS. For these experiments, 7 μM Cp183 dimer were used. The HAP12 ratio was calculated by accounting for HAP12’s racemic mixture, half of which is inactive, and that each dimer forms two HAP binding sites. Therefore, we used 14 μM HAP12 as a sub-saturating concentration, 28 μM as saturating, and 56 μM as super-saturating. These concentrations are well above the dissociation constant of HAP12 for capsid, so it is assumed all active HAP12 is bound (13). These concentrations corresponded to 1 active HAP12 molecules per dimer, 2 active HAP12 molecules per dimer, and 4 active HAP12 molecules per dimer, respectively.

Because different HAP12 regimens have different effects on capsid stability, we examined how they affected the kinetics of CTD exposure. The trypsin concentration for these experiments was chosen to avoid the much slower cleavage of the assembly domain (37). The loss of the parent Cp183 was expected to yield first order kinetics. Cp183 and cleavage products were visualized by SDS-PAGE and normalized to the total optical density of a given lane. For all nontreated and HAP12-treated data sets, a single exponential fit showed systematic differences in the rates of proteolysis. Data were well fit by two first order decays:

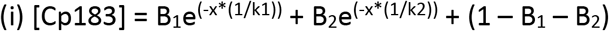

Coefficients B_1_ and B_2_ are the fraction of Cp183 in each of two populations of CTDs (Table 1) with k1 and k2 as the respective rate constants. A population of uncleaved CTD is explicit in the final term of this equation. From HAP12-free to saturating HAP12, the fast-cleaving CTDs of empty capsids accounted for 33% ± 5% of the Cp183 with a half-life of 0.4 ± 0.03 min. For the slow-cleaving population, the proteolysis accounted for 51% ± 8% with a half-life of 7 ± 2 min. At super-stochiometric HAP12, 74% of CTDs were rapidly cleaved with a half-life of 0.47 min; the remaining 25% had a half-life of 2.40 min.

**Table 1.** Curve fitting of CTD proteolysis data as a two first order decays.

| Sample | Population 1<br>(mole fraction) | Population 1<br>$t_{1/2}$ (min) | Population 2<br>(mole fraction) | Population 2<br>$t_{1/2}$ (min) | RMSD |
| --- | --- | --- | --- | --- | --- |
| Empty Capsid | 0.37 | 0.42 | 0.43 | 6.58 | 0.0065 |
| Empty + Sub-saturating HAP12 | 0.35 | 0.35 | 0.50 | 9.37 | 0.0105 |
| Empty + Saturating HAP12 | 0.28 | 0.40 | 0.60 | 5.51 | 0.0132 |
| Empty + Super-saturating HAP12 | 0.74 | 0.47 | 0.25 | 2.40 | 0.0033 |
| pgRNA-filled Capsid | 0.21 | 0.36 | 0.65 | 9.83 | 0.0101 |
| pgRNA Capsid + Super-saturating HAP12 | 0.24 | 0.29 | 0.72 | 7.73 | 0.0099 |
| E. coli RNA-filled Capsid | 0.24 | 0.28 | 0.12 | 13.97 | 0.0041 |
| E. coli RNA Capsid + Super-saturating HAP12 | 0.20 | 0.05 | 0.53 | 15.51 | 0.0068 |

Cp183 cleavage is in terms of populations cleaved rapidly, cleaved slowly, and uncleaved. The mole fractions of these three populations sum to 1.0.

The cleavage kinetics for drug-free to saturated HAP12 datasets were very similar. It is not until there was super-saturating HAP12 that we observed significant changes. Our data showed that HAP12 does not change CTD exposure in empty capsids until HAP12 reaches super-stoichiometric concentrations where it can induce capsid deformation that would expose more CTDs. Another issue to consider is whether the T=4 symmetry would modulate cleavage kinetics, i.e. would rates correspond to the A, B, C, and D subunits. The 33% average coefficient for the fast cleavage rate is far enough from the value of 25% predicted by quasi-equivalence, that quasi-equivalence seems an unlikely explanation. We suggest that cleavage rates may be affected by progressive charge changes within the capsid during proteolysis as well as capsid heterogeneity along with quasi-equivalence.

The pattern of CTD cleavage is also affected by capsid content. For HAP12-free empty capsids, proteolysis with trypsin yielded three doublets of cleaved Cp183 and a faint seventh band by SDS-PAGE (Figure 2A). For empty capsids treated with super-saturating HAP12, all Cp was proteolyzed to the lowest molecular weight band (Figure 2A). Reaction mixtures were further analyzed by LC-MS to identify SDS-PAGE bands. The highest molecular weight band corresponded to Cp183, and the lowest molecular weight band to the N-terminal 150 residues of Cp (Figure 2A). Other LC-MS data indicated cleavage after residue 175, 157, 152, and 151 (Figure 2A). These are consistent with previous CTD proteolysis studies (28).

**Figure 2.**
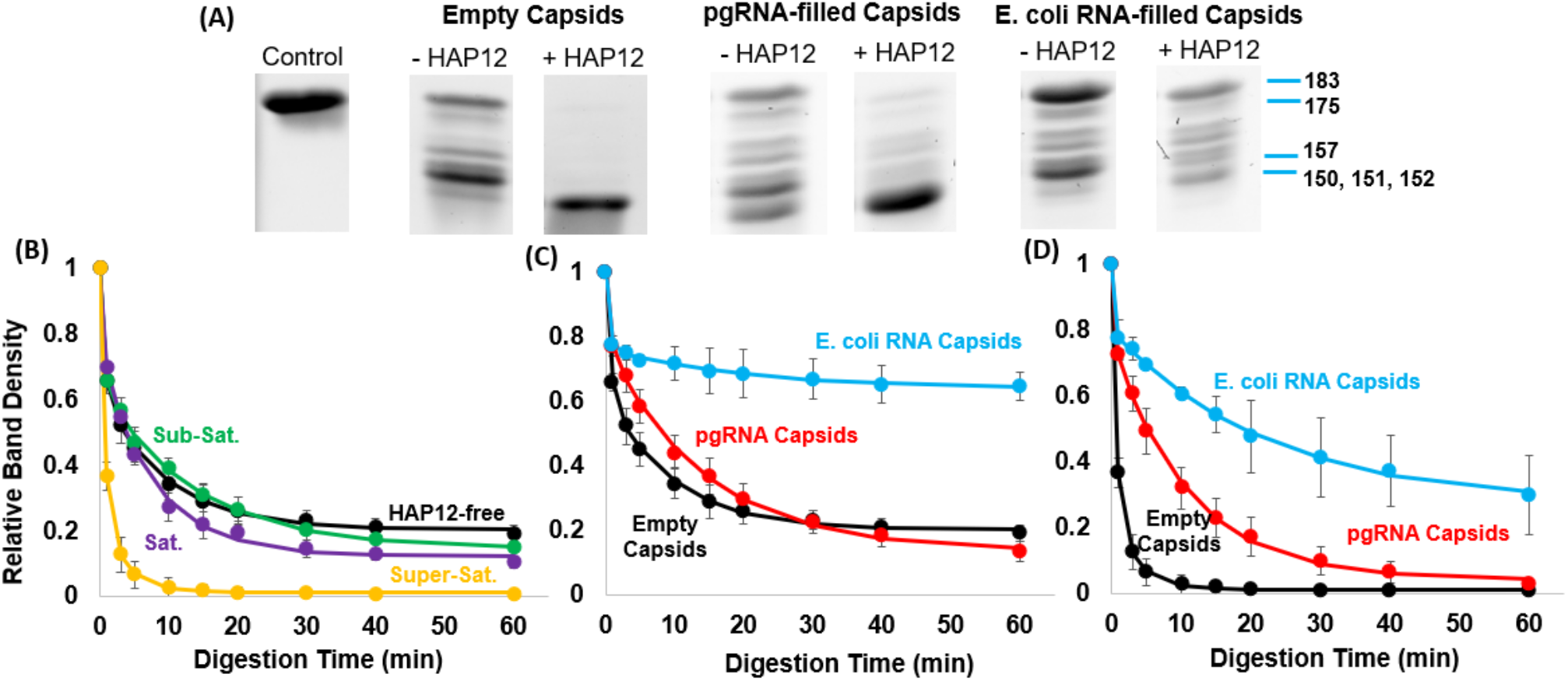
Excess HAP12 changed the proteolysis pattern and led to faster and more complete proteolysis of CTDs. CTD exposure was measured as a function of trypsin digestion. Empty, pgRNA-filled, and E. coli RNA-filled capsids were treated with HAP12 and then exposed to trypsin at room temperature. Capsid and trypsin concentrations were 7 μM and 0.04 μM, respectively. Reactions were quenched by acidification and analyzed by SDS-PAGE and LC-MS. (A) SDS-PAGE of proteolyzed, drug-free capsids (-HAP12) and capsids treated with super-saturating concentration of HAP12 (+HAP12) capsids; these samples were digested for 60 min. Prominent bands, identified by LC-MS, are labeled on the rightmost image. Untreated capsids retained substantive fractions of intact Cp183. With excess HAP12, the CTDs of empty and pgRNA-filled capsids were almost completely digested; E. coli RNA-filled capsids exhibited about equal distribution of different proteolyzed bands. (B, C, D) Time courses of cleavage at (B) varying HAP12 concentrations, (C) varying capsid content (empty, pgRNA-filled, and E. coli RNA-filled) without HAP12, and (D) varying capsid content with super-saturating HAP12. The relative band density is defined as the optical density of undigested Cp183 divided by the total optical density of the lane.

We anticipated that pgRNA-filled capsids would be more resistant to proteolysis than empty capsids since electrostatic interactions between the RNA and CTDs would keep these flexible peptides inside (18). We also predicted that pgRNA capsids would become more susceptible to proteolysis after HAP12 treatment, based on our proteolysis results with empty capsids (Figure 2A). For these experiments, purified Cp183 dimers were assembled with *in vitro* transcribed pgRNA to produce RNA-filled capsids. In the absence of HAP12, pgRNA-filled capsids were slightly slower to show signs of proteolysis. However, the pgRNA-filled capsids yielded a cleavage pattern distinctly different from those of empty capsids (Figure 2A). An important difference was that pgRNA-filled capsids had a prominent band for cleavage to residue 150, almost absent in empty capsids, suggesting that for at least a subset of subunits the CTDs were more easily digested to the junction with the assembly domain. The pgRNA capsids also had a different relative distribution of the cleavage doublets. In the presence of super-saturating HAP12, pgRNA-filled capsids were more resistant to proteolysis than empty capsids, retaining small amounts of partially cleaved Cp183 after one hour, but more sensitive to proteolysis than pgRNA-filled capsids without HAP (Figure 2A).

As an alternative to *in vitro* assembled pgRNA-filled capsids, we also tested E. coli RNA-filled capsids. When expressed in E. coli, Cp183 dimers assemble around non-specific E. coli RNA (20, 38, 39). For E. coli RNA-filled capsids in the absence of HAP12, the most prominent band after trypsin treatment remained intact Cp183, showing that these capsids were more resistant to proteolysis than empty and pgRNA-filled capsids (Figure 2A). Digestion of E. coli RNA-filled capsids with super-saturating HAP12 produced almost equal distributions of each proteolysis product. The E. coli RNA capsids were much more resistant to trypsin than pgRNA-filled capsids. E. coli RNA-filled capsids had a larger population of uncleaved CTDs at one hour (64% and 27%, respectively) compared to those of pgRNA-filled capsids (24% and 4%, respectively) (Table 1). However, with no HAP12 or excess HAP12, both RNA capsids had a similar fast-cleaving population of 22% ± 2%, which is close to the 25% predicted for quasi-equivalence. The data suggests that a quasi-equivalent class of monomers are less protected by the internal RNA and more accessible for proteolysis.

The difference in cleavage pattern and extent suggested fundamental mechanistic differences in CTD exposure. To compare the differences, we measured the rates of loss of Cp183 for empty, pgRNA, and E. coli RNA capsids, with and without super-saturating concentrations of HAP12. On a relative scale, depending on the presence of HAP12, empty capsids exhibited the fastest hydrolysis, pgRNA-filled capsids showed an intermediate rate, and E. coli RNA-filled capsids had the slowest hydrolysis (Figure 2B, 2C, & 2D). In the absence of HAP12, digestion appeared to stall for all three capsid types tested. In the presence of super-saturating concentrations of HAP12, empty and pgRNA capsids lost 100% of the initial Cp183 (Figure 2D and Table 1). For E. coli RNA-filled capsids, CTDs were hydrolyzed slightly faster than in the absence of HAP12 but was still the slowest of the HAP-bound capsids; also, it was not clear whether digestion would proceed to completion with longer incubation. For both untreated and treated conditions, our data demonstrated that the RNA content of a capsid can modulate the kinetics of CTD exposure. However, the RNA-CTD interaction does not completely trap CTDs within the capsid and prevent exposure.

### HAP12-treatment causes capsid deformation

Our proteolysis data indicated that HAP12 caused local structural changes that increased CTD exposure. To determine how these observations correlate with global changes in capsid morphology, we treated empty capsids with saturating HAP12 and examined them by negative stain transmission electron microscopy (TEM). Previous work with HAP-TAMRA and Cp149 capsids showed that HAPs caused deformation of normal capsids (15); similar behavior was seen with a dibenzothazepine CpAM (40). We anticipated that HAP12-treatment of empty Cp183 capsids would also lead to capsid deformation. Capsid samples were stained with ammonium molybdate and trehalose, where trehalose was used to maintain the 3D structure of capsids by minimizing the sample flattening that occurs when drying grids. In representative micrographs, HAP-treated capsids show a relatively small number of divergences from control particles (Figure 3A and 3B). 2D class averages tell a more detailed story. Class averages of untreated capsids showed nearly circular particle projections (Figure 3C); this morphology is also seen in cryo-EM (15). Class averages of HAP12-treated capsids displayed distinctly deformed structures; capsids were faceted, elongated, and broken (Figure 3D). Faceted and elongated capsids were the most common. The first ten classes accounted for 86% of the images. It was not clear how their global structural changes affected local regions of the capsid and led to greater CTD mobility. The broken capsid classes, which only accounted for 4% of the images, showed unambiguously how internal CTDs could be probed by proteolysis. It is possible that local changes could lead to greater CTD exposure or that HAP treatment promoted transient ruptures. In addition, we noted that the asymmetry of these classes could explain why the populations from our proteolysis curve-fitting (Table 1) was not consistent with quasi-equivalence.

**Figure 3.**
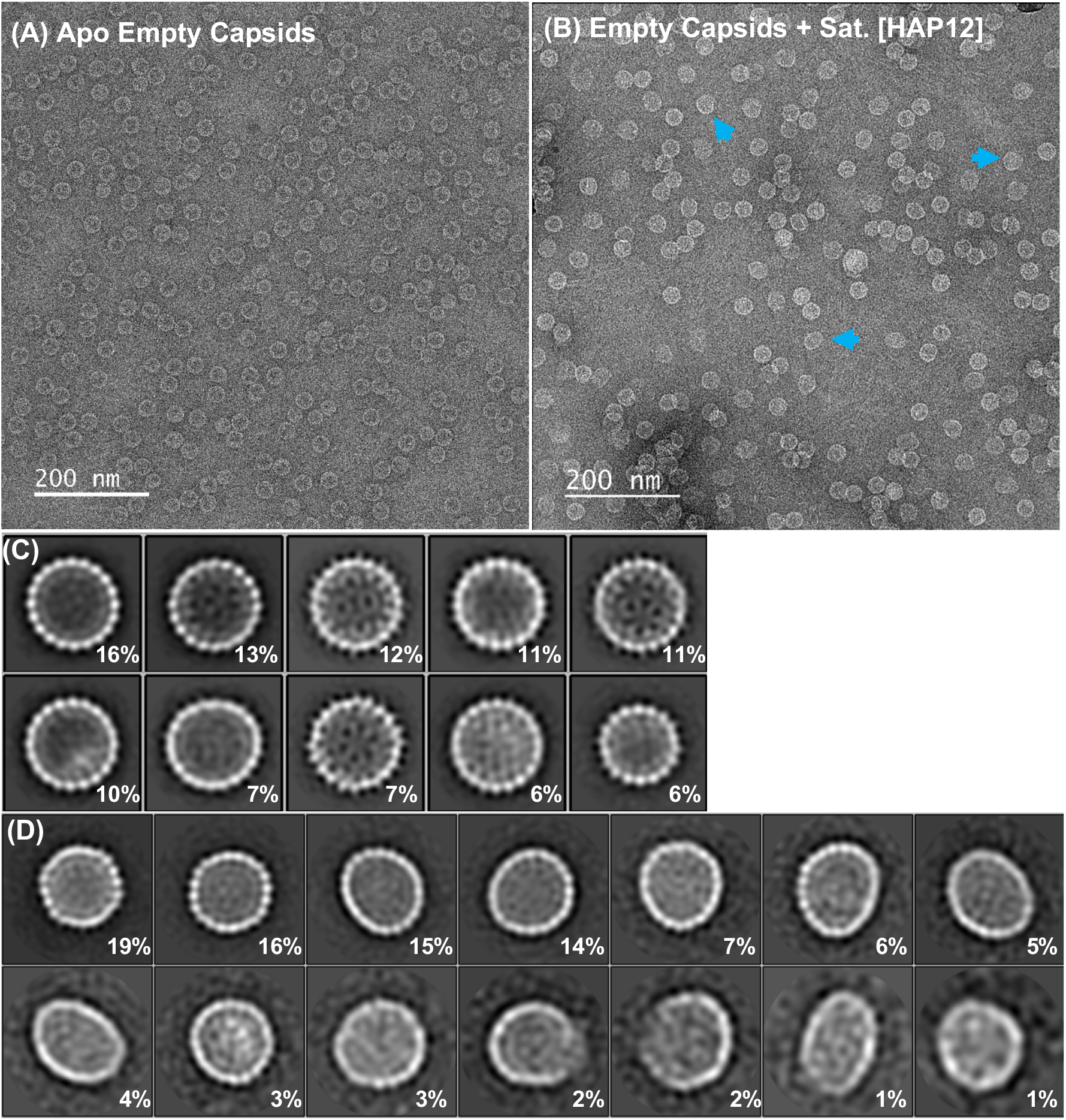
HAP12-treatment causes capsid deformation. Untreated (A,C) and treated (B,D) empty capsids were compared by negative stain EM. (A) A TEM of capsids with no HAP12 shows a narrow range of diversity. (B) A TEM of capsids in saturating HAP12 shows that damage to capsids is evident even without averaging. Blue arrows indicated deformed capsids. Samples were stained with ammonium molybdate with trehalose added to minimize sample flattening and distortion due to drying. (C) Class averages of 5,307 nontreated-capsids from negative stained TEM show a circular cross section. (D) Class averages of 1,160 capsids treated for 2 hours with saturating HAP12, at 25°C, show elliptical, faceted, and broken morphology; capsid deformation by HAP12 is nearly universal but irregular. Population of each class is shown in the bottom right corner.

### HAP12 with Impβ led to disruption of empty capsids, but E. coli RNA-filled capsids appeared unperturbed

The proteolysis data indicated that high HAP12 concentrations led to more CTD exposure in terms of the fraction of CTDs exposed, the length of the exposed peptide, and the rate of exposure. TEM data confirmed that HAP12 treatment causes capsid deformation, which correlates with greater CTD accessibility. This led us to question whether HAP12-treatment increased capsid binding to a biologically relevant ligand, the nuclear transport protein Impβ, and whether the stress of two different capsid ligands, HAP12 and Impβ, changed capsid integrity. Initially, we tested and confirmed Impβ binding to empty capsids under HAP12-free and HAP12-treated conditions (Supp. Figure 1A–C). To observe capsid morphology, untreated and HAP12-treated empty and E. coli RNA-filled capsids were mixed with Impβ, dialyzed into low salt, and observed via negative stain TEM. Control micrographs show that untreated capsids, empty and E. coli RNA-filled, appeared spherical and intact (Figure 4). Addition of Impβ had little effect on capsid morphology. We observed similar capsid morphology when sub-saturating HAP12 and Impβ were added to empty and E. coli RNA-filled capsids. However, we note that, at sub-saturating HAP12, the capsid border appeared uneven and thicker than those of non-drugged capsids. The thicker capsid shells may be due to bound Impβ. The observation of thicker capsid walls made us curious if sub-saturating HAP12 stabilized empty capsids, reduced internalization of Impβ, and led to more externally bound Impβ. To test this, 2D class averages were generated to determine the morphology of these capsids.

**Figure 4.**
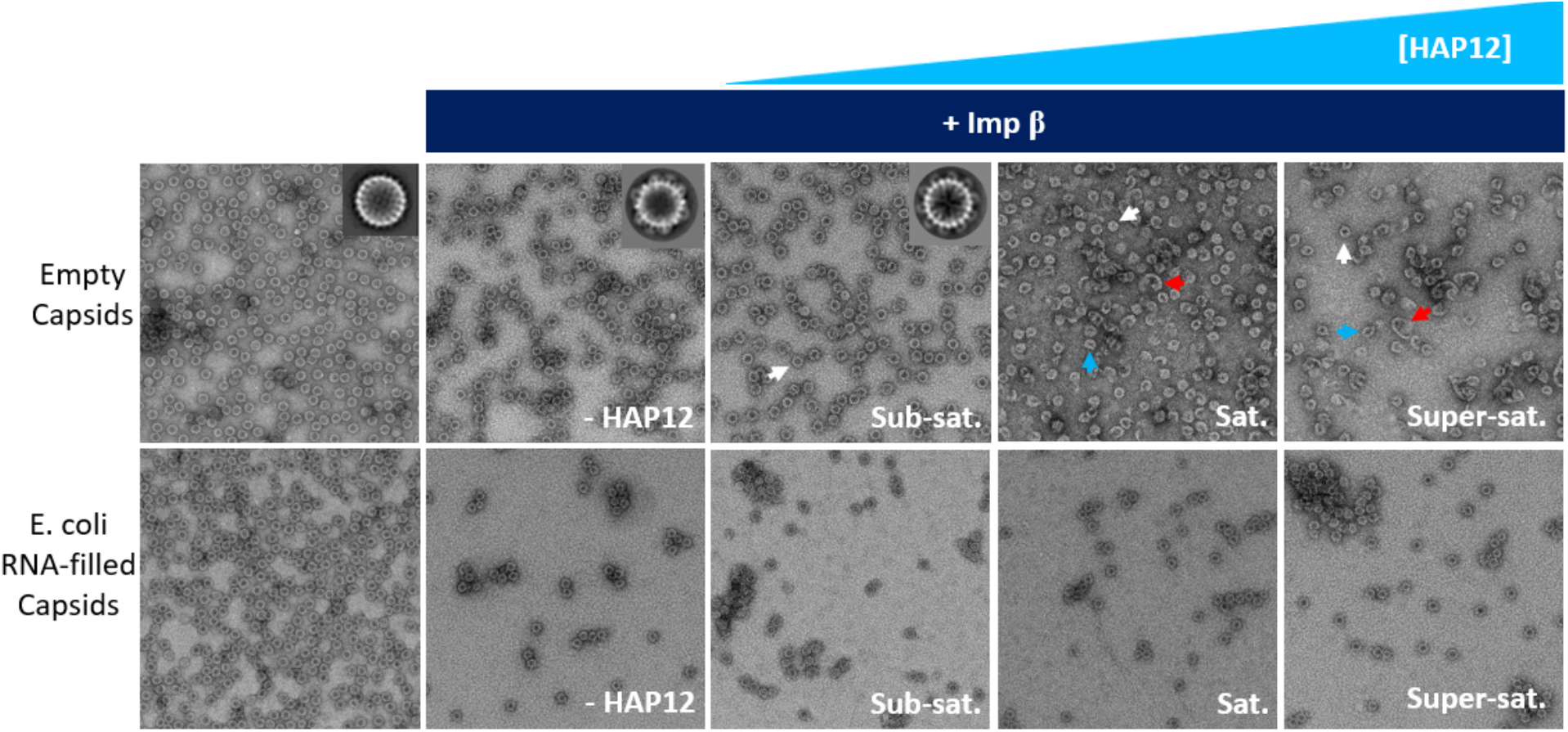
HAP12 with Impβ led to disruption of empty capsids. Empty and E. coli-RNA-filled capsids were mixed with Impβ so that there were 80 Impβ per capsid (Cp183 dimer and Impβ concentrations were 11.9 μM and 8 μM respectively), dialyzed into low salt (150 mM NaCl) to facilitate binding, and visualized by negative stain TEM. Relative HAP12 concentrations are shown in white. Arrows highlight capsids of specified morphology: white arrows – round, “normal”, capsids; blue arrows –deformed/intact capsids; red arrows – broken capsids/aberrant structures. Drug-free empty capsids appeared spherical and intact, which is reflected in its representative 2D particle projection (inset). After addition of Impβ, capsids with no HAP12 or sub-saturating HAP12 concentrations similarly appeared round and intact; each have representative cross-sections that show well-ordered, external Impβ density (insets and Supp. Figure 2). Introduction of Impβ and saturating or higher concentrations of HAP12 to empty capsids produced broken capsids and heterogenous complexes (see also Figure 5). E. coli RNA-Filled capsids remained morphologically unchanged even with treatment with high concentration of HAP12.

For empty capsids with Impβ, based on class averages, 48% of particles showed visible, externally bound Impβ, while 32% of particles showed no Impβ (Supp. Figure 2A). One class, accounting for 13% of particles, had density within the bounds of the capsid walls, suggesting internalized Impβ as observed in earlier studies (26). It is possible that in classes without evident Impβ, the importin adopted varying conformations so that the signal was blurred into background during averaging. With Impβ and sub-saturating HAP12, 90% of empty capsids showed unambiguous, externally bound Impβ (insets in Figure 4 and Supp. Figure 2B). This observation suggests that these Impβ adopted similar localizations. The Impβ density (“U” or “J” shaped) and arrangement were clear enough to allow us to determine which symmetry axis we were looking at, based on the number of external Impβ in each class. Ten external Impβ separated at the same intervals on the particle surface suggested that we were looking down a five-fold axis (Supp. Figure 2B). Eight external Impβ suggested that we were peering down a twofold axis. Two classes, accounting for 7% and 2% of classified particles, had strong internal density and two (17% and 10%) had internal “U” shaped features that were suggestive of internalized Impβ but could also be CTDs (some of these observations are addressed in the CDMS analysis below).

E. coli RNA-filled capsids appeared unperturbed and maintained normal capsid morphology even with excess HAP12 (Figure 4). 2D class averages of these samples confirmed that E. coli RNA capsids were able to maintain capsid geometry despite the presence of Impβ and super-stoichiometric CpAM (Supp. Figure 3B). It appears that the interaction between internal RNA and CTDs reduces all or most subunit modulation caused by HAP12-binding. However, we note that our proteolysis data still indicated super-saturating HAP12 changed CTD exposure (Figure 2 and Table 1).

When saturating or super-saturating HAP12 and Impβ are added, empty capsids became a heterogenous population of spherical capsids, deformed but intact capsids, ruptured capsids, and large Cp oligomers. To avoid the artifacts of negative stain TEM, we performed cryo-electron tomography to characterize the diverse population of deformed and disrupted particles. The tomograms showed new 3D structural features (Figure 5). Consistent with negative stain TEM, we observed a range of Cp oligomers. The largest oligomers had clear regions of hexagonal patterning. Not seen in negative stain images, we also observed areas of flat Cp sheets. Some sheets were relatively small and suggested a capsid that had unraveled in favor of planar geometry, leaving numerous gaps. Other oligomers were much larger than capsids, indicating that subunits, or oligomers of subunits, had been released and subsequently re-associated. The tomograms show how small molecules that impair Cp assembly also disrupt capsids and rearrange subunit geometry.

**Figure 5.**
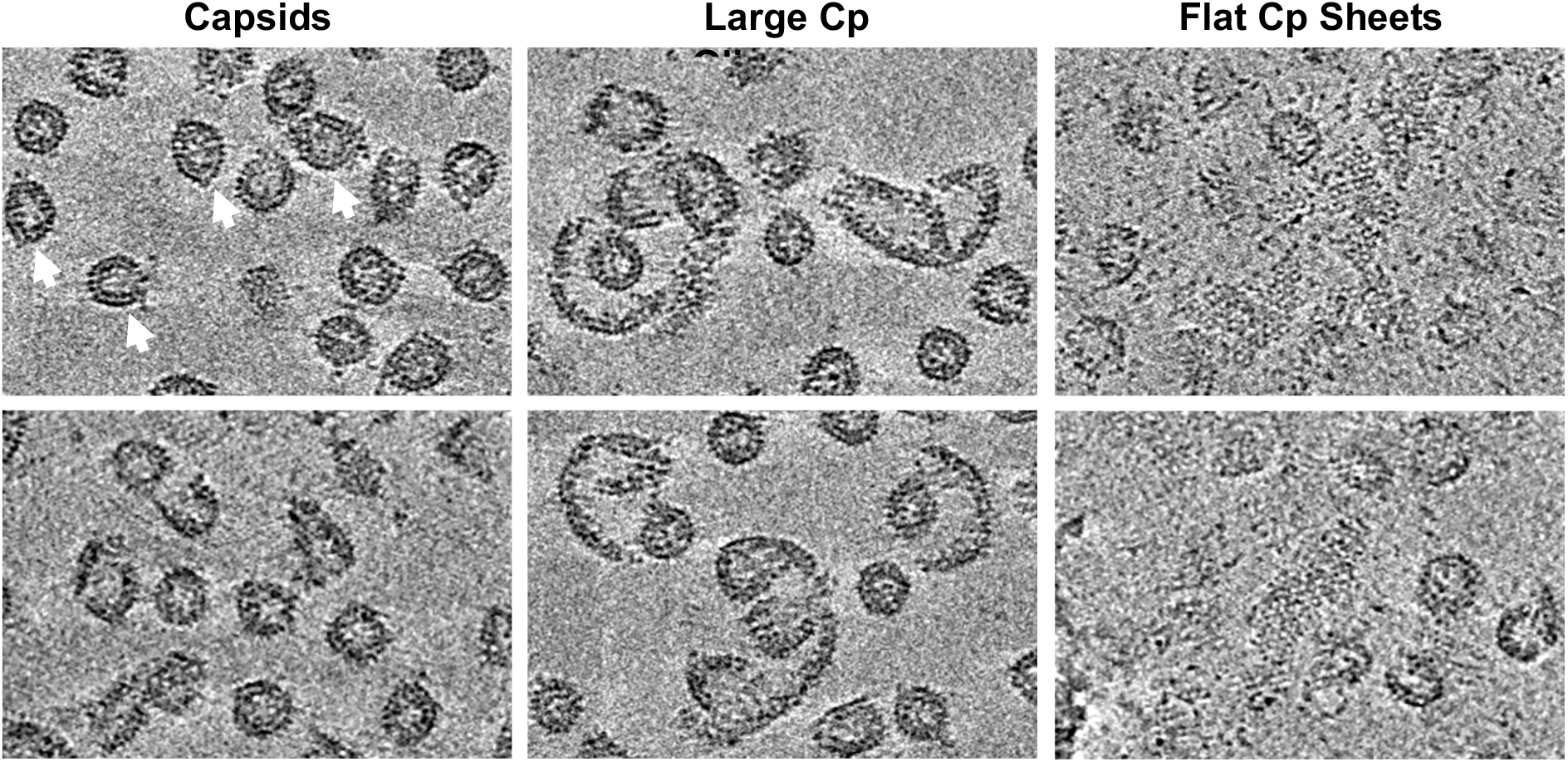
Cryo-electron tomographs of Impβ and HAP12-bound empty capsids revealed a diverse group of particles and Cp oligomers. Selected slices from tomograms of capsids treated with saturating HAP12 (for 2 hours) and Impβ (16 hours) suggest a progression of events. Capsid-sized objects, some of which are clearly distorted and broken (arrows), are the most common species (left column). Large Cp oligomers have many more dimers than a capsid and have a much larger radius of curvature (middle column). Flat Cp sheets are seen in some tomographic sections; these may result from a capsid unfurling or may result from free dimers self-assembling with planar geometry (right column).

### CDMS of empty capsids ± Impβ ± HAP12 showed shifts in mass and charge

The combination of HAP12 and Impβ led to structural and physical-chemical changes to empty capsids. This suggested that HAP12-induced capsid deformation could influence the number of Impβ binding to capsids. We predicted that saturating HAP12 treatment would allow more Impβ to bind to capsids compared to apo capsids and that greater Impβ binding would correlate with more capsid disruption. To measure the number of bound Impβ, capsid complexes were analyzed by charge detection mass spectrometry (CDMS). CDMS measures the mass-to-charge ratio (m/z) and charge (z) of each ion; these values are multiplied to give the mass of each ion, making it particularly powerful when the sample is heterogenous. The surface area of a particle dictates the number of charges on the ion; an approximation of the charge on a spherical water droplet, the Rayleigh limit, is a predictor for the number of charges on a spherical particles (41), while highly charged ions located above the limit are likely to have more extended or textured surfaces (42).

Empty capsids exhibited a narrow peak at 5.3 MDa, which is consistent with the mass expected for a T=4 capsid (5.05 MDa) plus about 1% for counterions, salts, and water that remained closely associated the particle (Figure 6A1). In the charge versus mass plot, where each point represents an individual ion, we see that empty capsids were closely distributed along the Rayleigh limit (Figure 6B1). The 3D heatmap gives a clearer and more quantitative view of the distribution of ions in terms of mass and charge (Figure 6C1).

**Figure 6.**
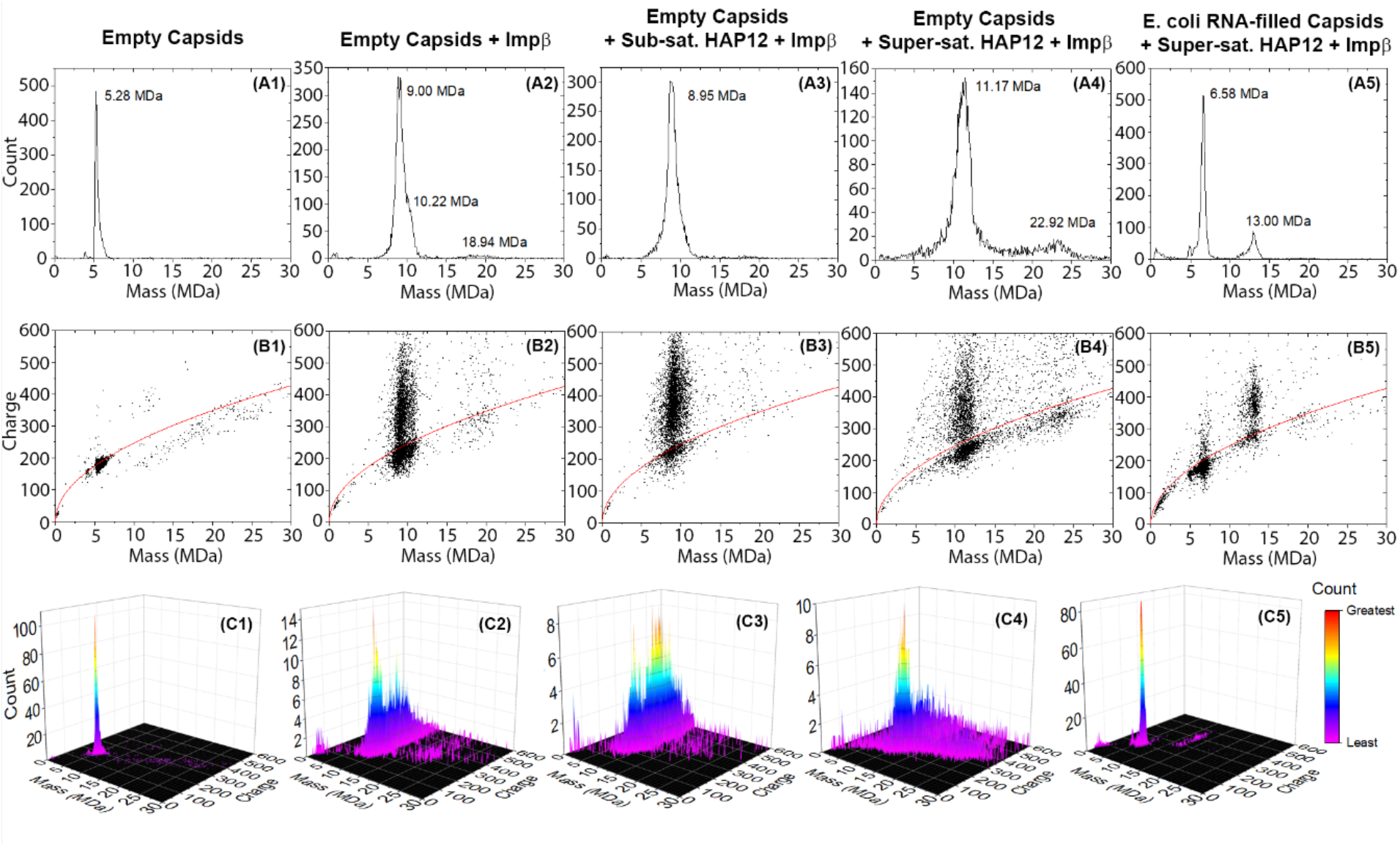
CDMS of empty capsids ± Impβ ± HAP12 showed shifts in mass and charge. Panels: A1-A5 mass spectra, B1-B5 charge vs. mass plots to identify individual ions (the Raleigh limit (red) shows the theoretical limit for the of charges on a spherical ion), and C1-C5 3D heat maps to identify distributions of populations. For empty capsids, mass shifts indicated Impβ binding to capsids. Most capsids showed two charged populations: a population near the Rayleigh limit, suggesting spherical particles, and a second, higher charged plume, suggesting extended or highly textured structures. Columns 1-4 show how empty capsids change mass and charge distributions in response to Impβ and/or HAP12. E. coli RNA-filled capsids (column 5) ± Impβ ± HAP12 exhibited no significant changes in their mass spectra, charge vs. mass plots, or 3D heatmaps.

When Impβ is added to empty capsids, the mass peak shifts to 9.0 MDa, indicating a modal average of about 38 Impβ molecules (97 KDa each) binding to each capsid (Figure 6A2). The charge versus mass plot shows two main populations of ions: a distribution near the Rayleigh limit and a vertical plume extending away from the limit (Figure 6B2). The charge plot suggests that this sample contained both compact spherical particles and extended or more textured particles. As the corresponding TEM data showed no broken capsids or other extended structures, we suggest that the highly charged distribution was attributed to Impβ-decorated capsids, which would be highly textured and could accept more charge. The 3D heatmap revealed that most of the ions were spherical (Figure 6C2). When sub-saturating HAP12 and Impβ are added, a similar mass was detected at 8.95 MDa, also corresponding to 38 Impβ per capsid (Figure 6A3). However, unlike the sample without HAP12, the 3D heatmap indicated that sub-saturating HAP12 led to more ions in the plume than in the cluster (Figure 6C3). This observation and the previous 2D class averages indicated to us that adding sub-stoichiometric HAP12 to empty capsids led to more externally bound Impβ. As a working hypothesis, in the absence of HAP12 and when a CTD binds Impβ, the capsid may internalize the host protein to relieve some of the mechanical strain on the capsid. However, with sub-saturating HAP12, CpAMs can strengthen Cp-Cp contacts and consequently suppress transient ruptures so that bound Impβ remains outside.

Super-saturating HAP12 and Impβ led to a shift in mass to 11.2 MDa, corresponding to approximately 64 Impβ per capsid (Figure 6A4). We also observed presence of many small and large ions, suggesting that some capsids had fragmented and some Cp183 dimers had coalesced into larger complexes (Figure 4 and Figure 6A4). This is consistent with tomographic data showing loosely connected sheets of protein and free subunits rearranging to create new structures (Figure 5). Of note, we observed a small broad peak at 22.92 MDa, which may be a dimer of two capsids; examination of the other mass spectra showed the same double capsid peak. Most of the ions attributed to capsids with Impβ and super-saturating HAP12 were near the Rayleigh limit, suggesting that most had a “spherical” morphology (Figure 6B4 and 6C4). The corresponding TEM images showed many “normal-sized” particles and a heterogeneous population of large oligomers (Figure 4). We also note that addition of saturating HAP12 produced similar mass and charge results as seen in the super-saturating HAP12 samples (Supp. Figure 4A5, 4B5, and 4C5).

While empty capsids showed evidence of CTD exposure and capsid deformation, at super-saturating HAP12, CDMS of E. coli RNA-filled capsids exhibited no mass shifts (Figure 6A5). Independent of all HAP12 concentrations, the peak attributed to E. coli RNA-filled capsids ranged from ~6.5-6.9 MDa, indicating little to no Impβ binding (Figure 6A5 and Supp. Figure 4A1–C4). This ~1.4 MDa mass difference compared to empty capsid is attributable to about 4,200 nucleotides of RNA. Most of the ions were near the Rayleigh limit, suggesting intact round capsids (Figure 6B5 and Supp. Figure 4B1–B4).

## DISCUSSION

Our study shows that exogenous molecules can work synergistically to disrupt HBV capsids. HAP12 globally disrupts capsid geometry and also modulates the accessibility of CTDs. CTD mobility and capsid stability are differentially modulated, depending on saturation of HAP sites. At low HAP12 saturation, CTD exposure remained unchanged compared to unmodified capsid, but capsids appeared to be more stable. At saturating or excess HAP12, CTD accessibility to protease increases, and capsids become disrupted with addition of Impβ’s. Additionally, encapsidated RNA modulated CTD exposure and increased capsid stability. E. coli RNA-filled capsids did not experience significant HAP-induced changes to their capsid geometry, as was observed with empty capsids. The RNA content of these capsids appeared to protect them against disruption. Though we were unable to test the sensitivity of a mature dsDNA-filled capsid to CpAMs and Impβ, the predicted and observed fragility of these particles (43, 44), led us to predict that they will be about as sensitive to disruption as empty particles.

This work shows that Cp-Cp interactions and consequently capsid stability can be exploited by small molecules. Like other CpAMs, HAPs exert seemingly paradoxical effects when bound to HBV capsids (16, 17). These small molecules increase the association energy of interdimer contacts (13, 15). However, by locally distorting the interdimer interface, HAPs can disturb global icosahedral geometry (15–17, 40, 45, 46). As more HAPs bind, a global cascading effect on capsid structure leads to deformation and eventually disruption (14–17, 40). HAP12 concentration also differentially affects CTD and capsid dynamics. In a T=4 capsid, there are A, B, C, and D pockets, each capped by a neighboring subunit. HAPs preferentially bind B and C sites; A and D sites are sterically hindered (see Figure 3a in reference (40)). In sub-saturating conditions, when HAP fills B and C sites, capsids are stabilized (13). We observe that when there is enough HAP to fill all four classes of site, capsids are destabilized because HAPs disrupt icosahedral geometry (14, 40). We showed that untreated or sub-saturating HAP12 conditions did not increase CTD sensitivity to protease activity in empty capsids. CTD accessibility only increases when HAPs reach a concentration threshold of saturating or higher. Therefore, Cp-Cp interactions experience a local stabilizing effect and global destabilizing effects from HAPs, depending on concentration, and CTD mobility is not directly affected by CpAMs.

The stabilizing and destabilizing effects of CpAMs can be exacerbated by Impβ. At sub-saturating HAP12, empty capsids do not show deformation but exhibit more externally bound Impβ than seen with undrugged empty capsids. This assertion is based on CDMS data showing that the +HAP12 +Impβ empty particles have a larger population of ions with charge in excess of the Rayleigh limit than the -HAP12 particles, suggesting that the +HAP12 particles have more textured surfaces (Figure 6C2 and 6C3). This observation is consistent with a sub-saturating HAP12, bound at a subset of pockets, that strengthens dimer-dimer interactions to prevent capsids from transiently breaking and internalizing bound Impβ’s (26). In addition, externally bound importins may further stabilize the capsids by neutralizing some of the positive charges at CTD clusters around the quasi-six-fold vertices (18). When we add saturating or super-saturating HAP12 along with Impβ, empty capsids deform, disrupt, and form Cp oligomers. Even though CpAMs strengthen interdimer contacts, it is not enough to offset the perturbance to capsid geometry which results in rearrangement of subunits. In addition to the effect of the CpAMs, the Impβ binding site extends beyond the CTD, which could exert a “pulling” force, partially unfold the dimer, or generally disturb the capsid’s quaternary structure (26). This interaction further compounds the mechanical strain on the capsid.

Nucleic acid also impacts CTD and capsid molecular motion. Internal RNA influences the organization of CTDs (18). Here we observed that differences in RNA content affect capsid behavior. In this study we observed that CTD exposure was slower with both RNA capsid types that were tested. Surprisingly, pgRNA-filled capsids were much more protease sensitive than E. coli RNA-filled capsids. Similarly, our data indicated that no Impβ bound to E. coli RNA-filled capsids (Figure 6A5, 6B5, and 6C5; Supp. Figure 4A1–C4), despite the presence of excess HAPs and CTD accessibility to proteases (Figure 2A–D and Table 1). This lack of binding to E. coli RNA-filled capsid was also observed with serine-arginine protein kinases (25). We surmise that electrostatic RNA-CTD interactions inhibit CTDs from externalizing to the capsid exterior. We also observed that E. coli RNA capsids do not experience detectable capsid deformation after treatment with excess HAP12 (Figure 4). We suggest that CTDs electrostatically crosslink dimers via their interaction with packaged nucleic acid to overcome capsid distortion.

Although RNA-filled capsids appeared more stable than empty capsids, their stabilities are further differentiated by their net internal charges. Our pgRNA-filled capsids have a net +200 charge: −3200 for the RNA and +3360 for the CTDs (+14 per Cp monomer from 16 arginines, one glutamate, and the C-terminus). Based on our CDMS data, the E. coli RNA-filled capsids used in these experiments have about 1.4 MDa of RNA, about 4,200 nucleotides from an undetermined number of polynucleotides, resulting in a net −840 charge (Figure 6A5, 6B5, and 6C5; Supp. Figure 4A1–C4). With or without excess HAP12, pgRNA capsids showed more cleavage than E. coli RNA capsids (Figure 2A–D and Table 1). The excess positive charge of pgRNA-filled capsids indicates that not all CTDs can be protected. Furthermore, the single long pgRNA may have limited access to CTDs (47); pgRNA is just long enough to coat the interior surface of the capsid (48). Conversely, the packaged E. coli RNA is present in shorter and longer sizes that can avoid the constraints of a single polynucleotide (19, 47, 49).

We propose that HAP12 and Impβ can strain a capsid and that above a threshold of strain, a ruptured capsid is more stable than an intact one – an accumulated strain model applied to icosahedral viruses (40, 50). We propose that the free energies of a HAP12 and Impβ-bound capsid in its (ii) strained and (iii) ruptured states are defined as:

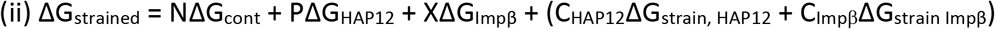

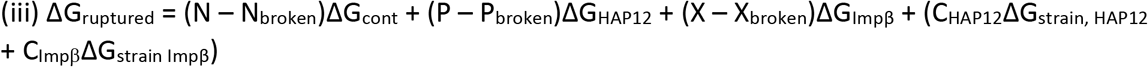

First consider an intact capsid (equation ii). N accounts for the 240 contacts made by the 120 dimers in a T=4 capsid. Based on structures of capsids with bound HAPs (15, 46), we assume that HAP12 binds to the 120 B and C sites in a T=4 capsid, noted as P_sites_ (15, 46); HAP bound to the 120 disfavored (and destabilizing) A and D sites is accounted for in ΔG_strain, HAP12_. The values for ΔG_cont_ and ΔG_HAP12_, which are −3.1 kcal*mol^−1^ and −1.9 kcal*mol^−1^ per contact, respectively. ΔG_Impβ_ is the increment that a bound Impβ stabilizes a capsid, presumably by neutralizing some electrostatic repulsions of the CTDs; empty capsids disassemble and precipitate in the absence of excess ionic strength (28) or Impβ. X_Impβ_ is the number of bound Impβ molecules, less than or equal to 240. Finally, in an intact capsid, there are strain terms attributable to excess bound HAP (accounting for geometric effects and binding to disfavored sites) and Impβ.

A ruptured capsid loses stabilizing and destabilizing energy to reach a minimum (equation iii). N_broken_ represents contacts that are lost after capsids rupture. P_broken_ accounts for lost HAP12 molecules that were initially bound at interdimer interfaces but are released once their contacts break. X_broken_ represents Impβ-bound dimers that have disassociated after rupture. Finally, strain is lost: ΔG_strain, HAP12_ and ΔG_strain Impβ_ are influenced by capsid deformation, so they are multiplied by coefficients C_HAP12_ and C_Impβ_, respectively. These strain coefficients range from 0 - 1 and increase nonlinearly as the HAP12 and/or Impβ binding rises.

With sub-saturating HAP12, icosahedral capsids are favored: ΔG_strained_ < ΔG_ruptured_ (Figure 7B and 7D). However, at saturating and super-saturating HAP12, ΔG_strain,HAP12_ is large, and capsids undergo disruption to relax that strain (Figure 4; Figure 7E). An identical argument holds with high Impβ-binding. In conditions where rupture is favored the two C coefficients should be close to 0, so that C_HAP12_ΔG_strain, HAP12_ and C_Impβ_ΔG_strain Impβ_ are relaxed. Saturating HAP12 or above is required to disturb capsid geometry, and generate ΔGstrain,HAP12 (Figure 7B). Addition of Impβ compounds the capsid’s already strained state further drives the reaction to the ruptured state (Figure 7F).

**Figure 7.**
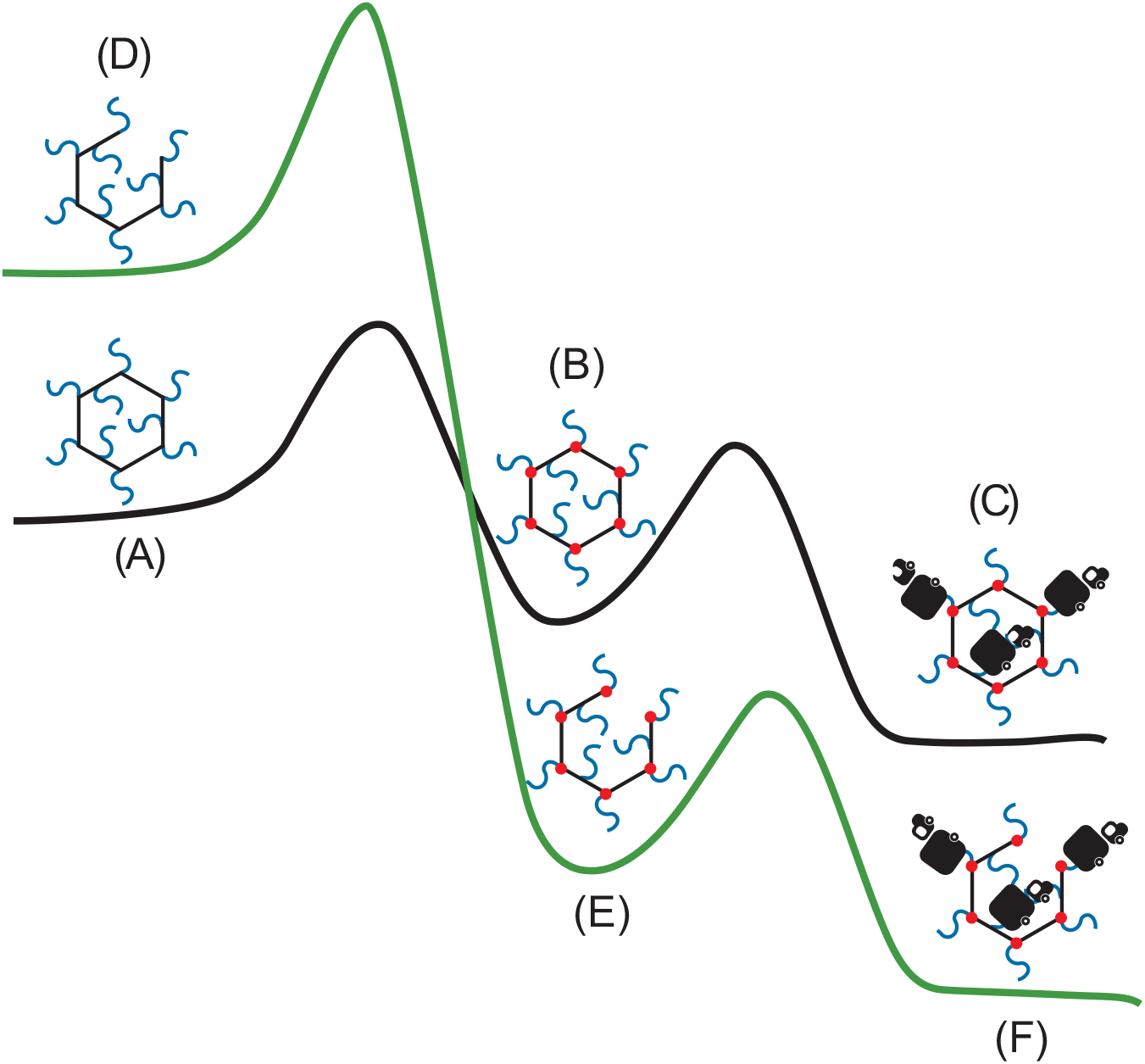
Free energy diagram of an empty capsid in its strained (A-C, black) and ruptured (D-F, green) states mediated by HAP12 and Impβ. Free energy schematics for a strained (A-C) or ruptured (D-F) empty capsid shown in black and green, respectively. Because it has fewer interdimer contacts, an untreated, ruptured capsid (D) is at a higher energy state than an intact, drug-free capsid (A). After enough HAP molecules (red dots) bind to cause capsid deformation, the strained, intact capsid (B) pays a global strain penalty and is at a higher energy level. By rupturing, a HAP-bound capsid relieves the global strain and achieves a lower energy state (E). Although Impβ-binding (black trucks) provides some stability to capsids, this interaction applies further mechanical strain on the capsid (C), which can be energetically compensated by rupturing (F).

This paper demonstrates and analyzes the dual antiviral activities of CpAMs and their synergism with host proteins. Molecules like CpAMs have been documented for their ability to modulate capsid assembly, but their ability to disrupt capsids have only been implicit until recently (13–15, 40, 46, 51). Here, we also show that host proteins, in particular Impβ, can contribute to capsid disruption. Destabilizing a capsid adds to the arsenal of mechanisms of action for small-molecule HBV therapeutics. During the HBV life cycle, empty and rcDNA-filled capsids interact with host importins and are trafficked to the nucleus (52). When capsids enter the nuclear pore complex, capsid protein will interact with Nup153, which may facilitate uncoating (32). Capsid disassembly at inopportune times can impair the virus’s ability to establish infection (16, 17). Furthermore, if rcDNA-containing capsids disassemble in the cytoplasm, the exposed viral DNA may be detected by the host’s innate immune system (e.g cGAS-STING (53)). Indeed, rcDNA capsids are fragile and may be particularly sensitive to disruption (43, 44, 54). Because Cp is involved throughout the HBV lifecycle, CpAMs continue to be a versatile and attractive candidate for developing an efficacious cure for chronic HBV infection.

## MATERIALS AND METHODS

### Protein Purification and Preparation

E. coli RNA-filled Cp183 capsids were expressed and purified as previously described (28). To isolate Cp183 for subsequent experiments, E. coli RNA-filled Cp183 capsids were dialyzed in 1.5 M guanidine hydrochloride, 0.5 M LiCl, 10 mM DTT, and 20 mM Tris-HCl pH 7.5 (disassembly buffer) at 4°C overnight. The sample was centrifuged at 7,000 g for 10 minutes to pellet the RNA. Cp183 dimers were purified from the supernatant using a Superose 6 column (GE) equilibrated in disassembly buffer. To assemble empty capsids, dimers were dialyzed in 0.45 M NaCl, 10 mM DTT, and 20 mM Tris-HCl pH 7.5 (assembly buffer) at 4°C overnight. To remove unassembled dimers, empty capsids were purified via SEC with a Superose 6 column that was equilibrated with assembly buffer. To prepare pre-genomic RNA (pgRNA)-filled Cp183 capsids, pgRNA was in vitro transcribed using the MegaScript kit (Thermo-Fisher) with plasmid 1135 (38), which was modified to have a T7 promoter. The modified plasmid 1135 was a gift from Dr. Dan Loeb. To assemble pgRNA-filled capsids, dimers and pgRNA were mixed in a 120:1 molar ratio and then dialyzed in 0.15 M NaCl, 10 mM DTT, and 20 mM Tris-HCl pH 7.5 at 4°C overnight. To produce E. coli RNA-filled capsids, stock E. coli RNA capsids were loaded onto a 10-40% continuous sucrose gradient and centrifuged at 40,000 rpm for 5 hours; the capsid band was extracted and dialyzed into assembly buffer. Impβ expression and purification protocols were adapted from Chen et al. (26).

### Treatment of HBV Capsids with the CpAM HAP12

For HAP12 treatment, empty and pgRNA-filled capsids were incubated with HAP12 for 2 hours away from light before they were used for experiments. To prepare drug stock samples, HAP12 was resuspended in 100% DMSO. HAP12 stocks contain a racemic mixture of 50% active and 50% inactive molecules; we always refer to HAP12 concentrations in terms of the whole racemic mixture. These concentrations correspond to the following HAP12 to dimer molar ratios: 2:1, 4:1, and 8:1, respectively. For example, at saturating concentration, there would be 4 HAP12 molecules for every 1 dimer, which corresponds to 1 active HAP12 molecule per pocket. For control samples, capsid samples were incubated with DMSO.

### Proteolysis of HBV Capsids

The limited proteolysis assay performed in this paper was adapted from a previously published work (28). Reaction samples with empty, pgRNA-filled, and E. coli RNA-filled capsids were diluted in 0.5 mM NaCl, 10 mM β-mercaptoethanol, and 80 mM Tris-HCl pH 7.5 and incubated with 0.04μM sequence grade modified trypsin (Promega) for specific times. Proteolysis was quenched with 4x Laemmli buffer and heating at 95°C for 6 minutes. Samples were then analyzed by SDS-PAGE on a 20% denaturing polyacrylamide gel, containing 0.4% (v/v) 2,2,2-trichloroethanol (TCE) to support direct fluorescent detection of protein samples (55). Gels were imaged with a ChemiDoc− (BioRad). Protein bands were quantified using the Fiji software (56). Each data point is the average of three independent trials with a corresponding error bar. Curve-fitting for CTD exposure kinetics was performed using the Solver function on Excel. For LC-MS, reactions were quenched by addition of glacial acetic acid.

### LC-MS of Proteolyzed Cp183

Two separate liquid chromatography-mass spectrometry (LC-MS) methods were used to characterize the peptides generated in the limited proteolysis experiments. The first focused on peptides with a molecular weight less than 3 kDa. Before analysis, samples with peptides were passed through a 3 kDa molecular weight cut off filter (PALL) to remove larger peptides and undigested protein. The final concentration of peptides was approximately 0.1 mg/mL. 1 μL of the peptide mixture was injected onto a Dionex 3000 nano-HPLC (Thermo-Fisher) coupled to a maXis Impact QTOF (Bruker Daltonics). Nano LC-MS/MS analysis was performed as as previously described. (57, 58). Briefly, LC was carried out using an Acclaim PepMap C18 reverse phase column (300μm x 5mm) using a slow gradient:0-2.5 min, 3% B; 2.5-20 min, 3-30% B; 20-23 min, 30-80% B; 23-25 mins, 80% B; 25-28 mins, 80-3% B; 28-30 mins, 3% B where solvent A = 0.1% formic acid (FA, Sigma) in water (Thermo-Fisher) and solvent B = 0.1% FA in acetonitrile (Thermo-Fisher). The peptides were analyzed using PeptideShaker (59) coupled to SearchGUI (60).

LC-MS of peptides greater than 3 kDa was performed using an Agilent 1290 UPLC series LC coupled to a micrOTOF (Bruker Daltonics). 10 μL of the peptide mixture at approximately 0.1 mg/mL were injected onto the LC. The LC was carried out using a Phenomonex Onyx Monolithic C18 reverse phase column (100 x 2mm) at 50° C with the flow rate of 400 μL/min. The following gradient was used: 1.0 min, 10% B; 1.0-8.0 min, 10-70% B; 8.0-8.5 min, 70-90% B; 8.5-9.0 min, 90-10% B; 9.0-10.0 min, 10% B solvent A = 0.1% formic acid (FA, Sigma) in water (Thermo-Fisher) and solvent B = 0.1% FA in acetonitrile (Thermo-Fisher). MS settings were as follows: nebulizer set to 5.0 bar, drying gas at 7.0 L/min, drying temperature at 200 °C, and capillary voltage at 4.5 kV. The capillary exit was set at 100V, skimmer 1 at 50V, hexapole 1 at 23V, hexapole RF at 300 Vpp, and skimmer 2 at 22V. Data analysis was carried out using Bruker DataAnalysis with MaximumEntropy.

### HBV Capsid and Importin β Binding Experiments

Impβ was passed through a Superose 6 column to exchange the buffer into 0.5 M NaCl, 10 mM DTT, and 20 mM Tris-HCl pH 7.5. Empty and E. coli RNA-filled capsids were mixed with Impβ in a 1:80 ratio. Then, these samples were dialyzed into 0.15 M NaCl, 10 mM DTT, and 20 mM Tris-HCl pH 7.5 at 4°C overnight to allow binding. The resulting complexes were analyzed via SEC, SDS-PAGE, CDMS, and TEM. For SEC, complexes were resolved using a Superose 6 column and the overnight dialysis buffer was the running buffer. SDS-PAGE was performed with a denaturing polyacrylamide gel (4% stacking and 16% resolving).

### Transmission Electron Microscopy

For negative stain TEM of empty capsid morphology, capsids were mixed with 6% (w/v) ammonium molybdate and 0.5% (w/v) Trehalose and applied to glow-discharged continuous carbon grids. For studies where sample flattening was not critical, samples were first applied to grids and then stained with 0.45% (w/v) uranyl formate or 2% (w/v) uranyl acetate. All grids were imaged using the JEOL 1400 FS microscope at 120 kV and at magnification 50,000x.

### Cryo-electron Tomography

To prepare cryo-EM specimens, a drop of 4 μL of sample mixture was applied to a glow-discharged 300-mesh Quantifoil^®^ R2/2 holey carbon grid. The grid was plunged into a liquid ethane bath cooled by liquid nitrogen, using a Thermo Fisher Scientific (TFS) Vitrobot Mark IV. The frozen hydrated cryo-EM grid was clipped into a cartridge and then transferred into a cassette before loading into a TFS 300-kV Titan Krios equipped with Gatan BioContinuum− energy filter, using a K3 direct electron detector camera. Data collection was set up using TFS Tomography software (v 4) under counting mode. The nominal magnification was 53,000x (equal to 1.7 Å per pixel), and the illumination had a dose rate of 0.7 e^−^/Å^2^. The zero-loss peak was aligned for each tilt-series, using an energy slit of 20 eV. Tilt series were acquired using a bidirectional scheme from −60° to 60° with tilt step at 2°. Therefore, the total accumulated dose is ~ 42.7 e^−^/Å^2^. Tilt series alignment, CTF correction, and tomogram reconstruction were performed using IMOD (v 4.9.12) (61). The final 3D reconstruction was generated with data binned at 4. Nonlinear anisotropic diffusion was applied to the final reconstruction to reduce noise (62).

### Charge-detection Mass Spectrometry (CDMS)

CDMS is a single particle technique in which the mass-to-charge ratio (*m/z*) and charge (*z*) of each ion are measured simultaneously. Multiplying the *m/z* by the charge gives the mass of an ion. Measurements are performed on many individual ions to generate a mass distribution. This allows molecular weight distributions to be measured for large (greater than 1 MDa) and heterogenous species. These species cannot usually be analyzed with traditional mass spectrometry methods that have an effective upper limit of about 1 MDa. In this work, a homebuilt CDMS instrument, described previously (63–65), was used. Briefly, the analyte is ionized by a nano-electrospray (Advion Triversa Nanomate). The ions enter the instrument through a metal capillary and pass three regions of differential pumping. They are then focused into a dual hemispherical deflection energy analyzer which transmits a narrow band of ion energies centered around the nominal ion energy of 100 eV/charge. The transmitted ions are focused into an electrostatic linear ion trap, where they are trapped when potential barriers are raised at both ends. As the ion oscillates back and forth in the trap, it passes through a conducting cylinder. The ion induces an equal but opposite charge on this cylinder; the signal from the induced charge is amplified, digitized and analyzed by fast Fourier Transforms (65). The fundamental frequency is proportional to the *m/z* of the ion, while the amplitude of the signal is proportional to the charge of the ion. All mass spectra were generated using a 100 ms trapping time. Typical spectra contain thousands of individual ion measurements and take 30-50 min to collect. Before CDMS analysis, empty and E. coli RNA-filled capsids samples were buffer exchanged into a volatile buffer, 150 mM ammonium formate pH 7.5 at 4°C. The CDMS data presented in this work includes mass spectra, charge versus mass plots, and 3D heatmaps. In addition, all charge versus mass plots contained Rayleigh limit curves. The Rayleigh limit is a model that predicts the surface charge of a water droplet of a certain mass (41, 66–68).

## ACKNOWLEDGEMENT

This research was supported by grants R01-AI144022 and R01-AI118933 from the NIH to AZ. We also acknowledge the Physical Biochemistry Instrumentation Facility and the IU Electron Microscopy Center, both which are supported the Indiana CTSI.

## Supplemental Figures

**Supplementary Figure 1.**
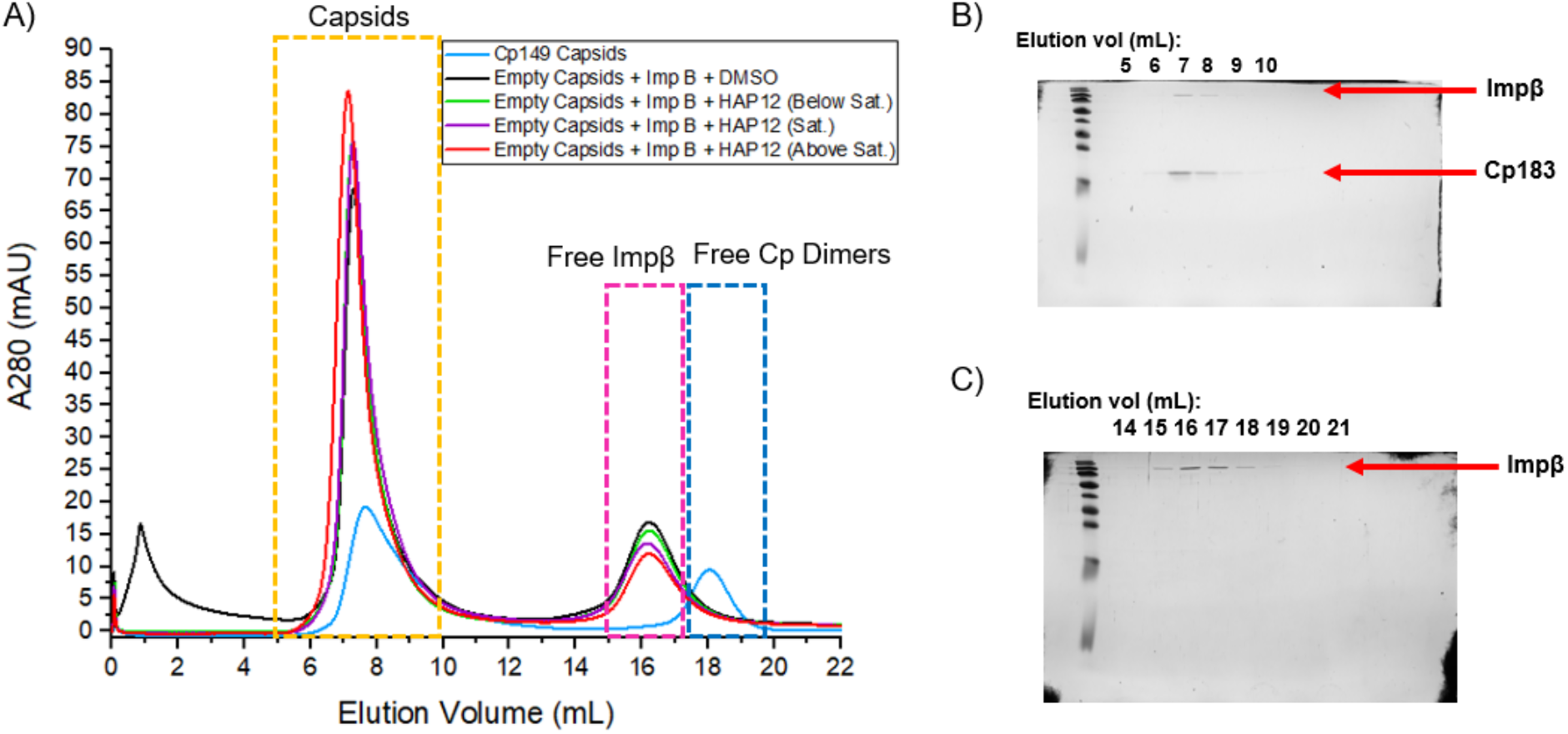
At super-sat. [HAP12], the capsid peak shifted to the left, which indicated elution of complexes larger than capsids. To resolve Impβ–bound capsids, 200 μL of each sample were injected onto a Superose 6 column. SEC fractions were collected and analyzed via SDS-PAGE. Panel A shows a compiled chromatograph of empty capsids ± Impβ ± varying HAP12 concentrations; a mixture of Cp149 capsids and dimers were run as a control. The dashed boxes designate at which volumes certain proteins elute at: yellow – capsids, pink – free Impβ, and blue – free dimers. Panels B-C show gel images of collected SEC fractions. Panel B shows co-elution of Cp183 and Impβ at 7-9 mL. Panel C shows elution of free Impβ at 15-19 mL.

**Supplementary Figure 2.**
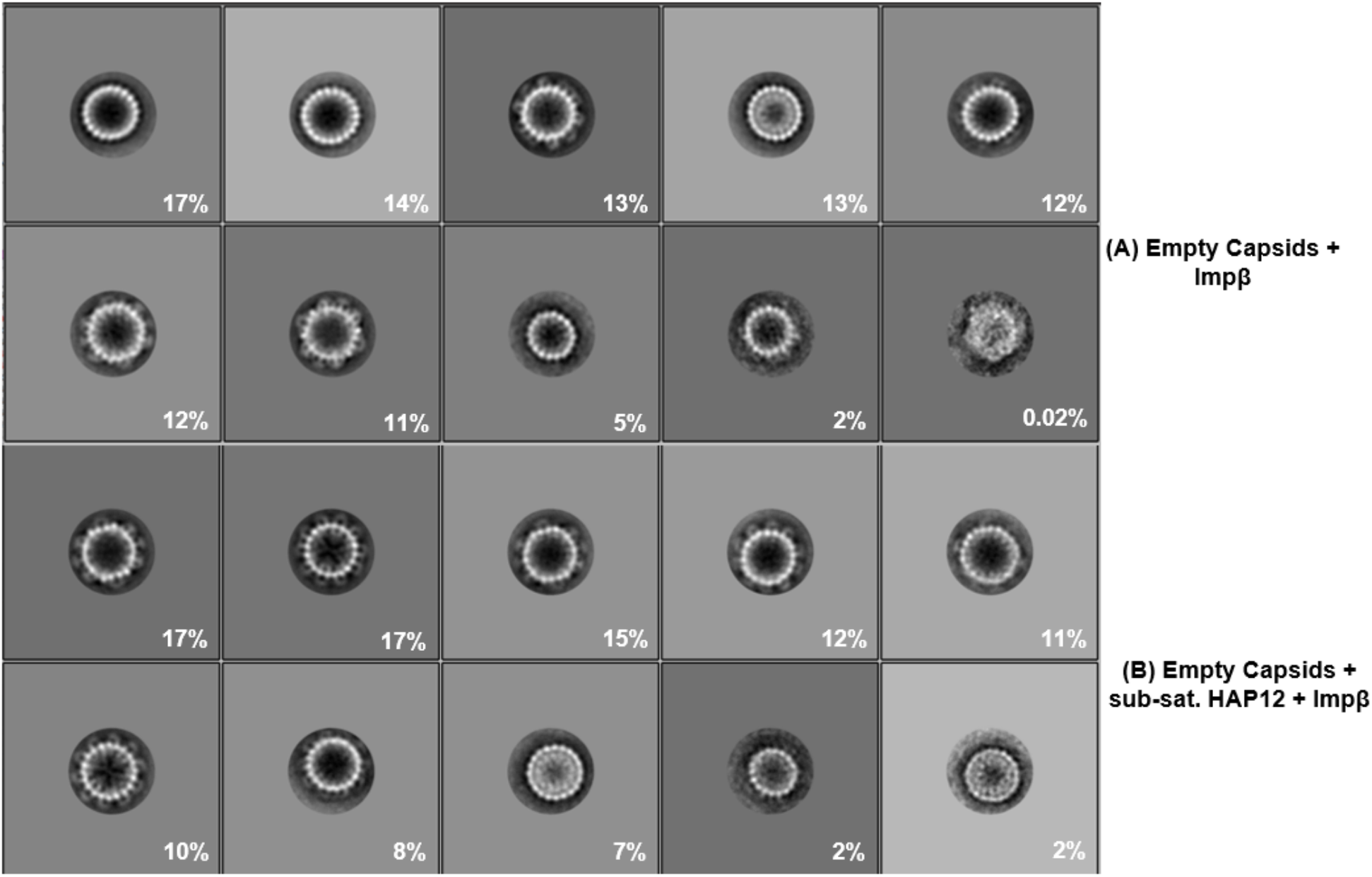
Empty capsids treated with sub-saturating HAP12 appeared to have more externally bound Impβ than drug-free capsids. Class averages of HAP-free empty capsids + Impβ (4,718 particles) and empty capsids + sub-saturating HAP12 + Impβ (5,118 particles) were compared for differences on capsid morphology (A and B). Empty capsids were prepared as previously stated in Figure 4. HAP12-free empty capsids showed four classes (48% of total particles) with visible, externally bound Impβ (A). However, capsids treated with sub-saturating HAP12 showed seven classes (90% of total particles) with external Impβ (B). These observations suggested that sub-stoichiometric HAP12 stabilized Cp-Cp interactions, preventing breakage of dimer-dimer contacts and internalization of Impβ.

**Supplementary Figure 3.**
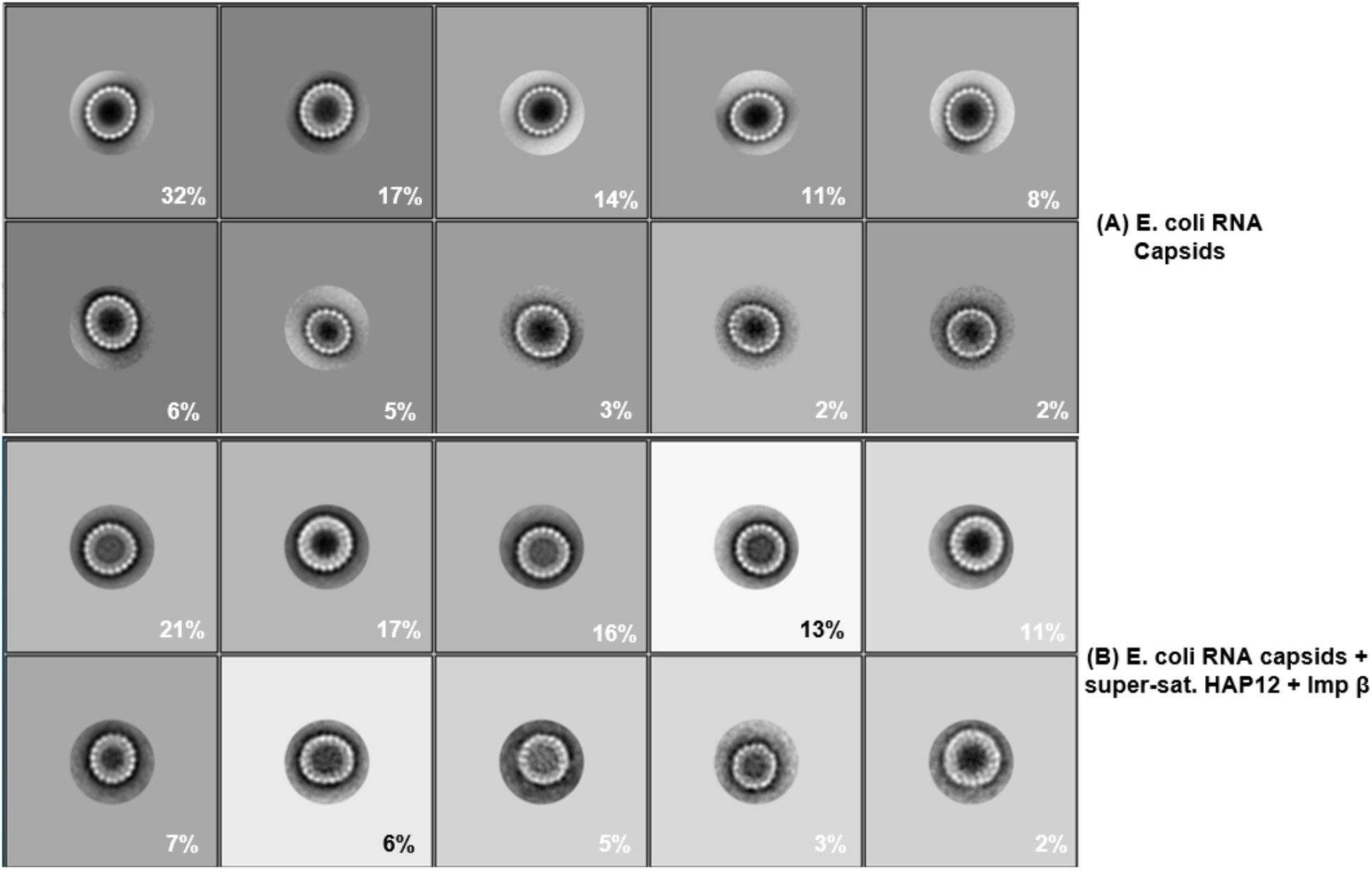
After treatment with super-saturating HAP12 and Impβ, E. coli RNA-filled capsids appeared intact and spherical. Class averages of E. coli RNA-filled capsids with no HAP12 (4,319 particles) or with super-saturating HAP12 + Impβ (4,247 particles) were compared for differences on capsid morphology (A and B, respectively). E. coli RNA capsids were prepared as previously stated in Figure 4. Untreated E. coli RNA capsids appeared intact and round (A). Similarly, E. coli RNA capsids treated with super-saturating HAP12 and Impβ also appeared to be intact and spherical, suggesting that internal RNA prevented capsid deformation and disruption (B).

**Supplementary Figure 4.**
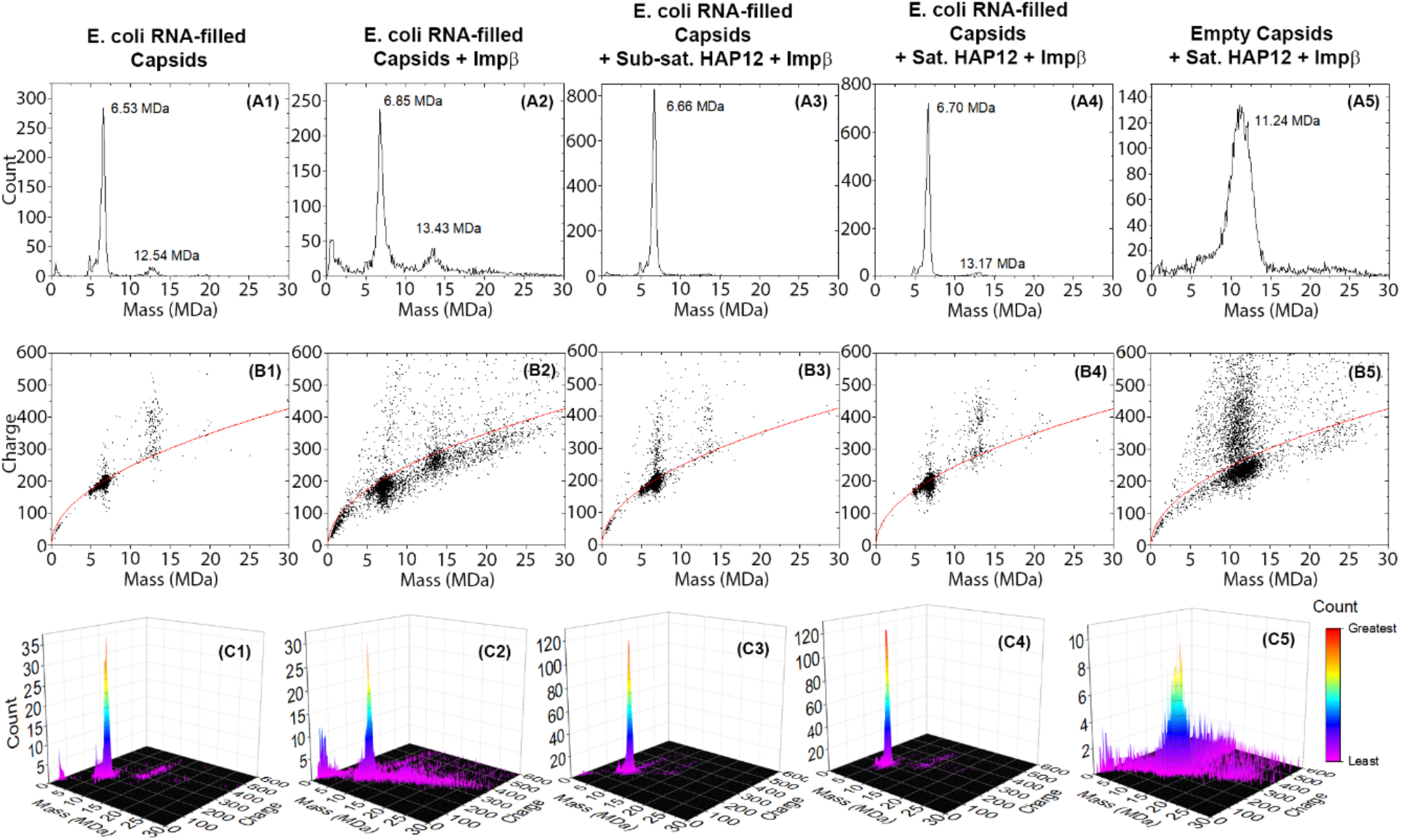
CDMS of E. coli RNA-filled capsids ± Impβ ± HAP12 showed no significant shifts in mass and charge. Panels: A1-A5 mass spectra, B1-B5 charge vs. mass plots with Rayleigh limit (red curve), and C1-C5 3D heatmaps. For E. coli RNA-filled capsids, no mass shifts were observed despite addition of Impβ or HAP12. For all E. coli RNA capsid samples, a concentration of ions was detected along the Rayleigh limit, suggesting spherical capsids. The 3D heatmaps show that E. coli RNA capsids fall into a single population of ions. Conversely, after addition of super-saturating HAP12 and Impβ, empty capsids undergo a mass shift and ion population changes that are similar to empty capsids treated with saturating HAP12 (Figure 6).

## References

1. Graber-Stiehl I. 2018. The silent epidemic killing more people than HIV, malaria or TB. Nature 564:24–26.

2. Polaris Observatory C. 2018. Global prevalence, treatment, and prevention of hepatitis B virus infection in 2016: a modelling study. Lancet Gastroenterol Hepatol 3:383–403.

3. Usha V, Nagraj M, Zhanying H, Haiqun B, Yanming D, Jin H, Jinhong C, Ju-Tao G. 2020. Targeting the multifunctional HBV core protein as a potential cure for chronic hepatitis B. Antiviral Research 182:104917.

4. Venkatakrishnan B, Zlotnick A. 2016. The Structural Biology of Hepatitis B Virus: Form and Function. Annual Reviews of Virology 3:429–451.

5. Beck J, Nassal M. 2007. Hepatitis B virus replication. World J Gastroenterol 13:48–64.

6. Prange R. 2012. Host factors involved in hepatitis B virus maturation, assembly, and egress. Med Microbiol Immunol 201:449–61.

7. Ceres P, Zlotnick A. 2002. Weak protein-protein interactions are sufficient to drive assembly of hepatitis B virus capsids. Biochemistry 41:11525–31.

8. Ning X, Nguyen D, Mentzer L, Adams C, Lee H, Ashley R, Hafenstein S, Hu J. 2011. Secretion of genome-free hepatitis B virus--single strand blocking model for virion morphogenesis of para-retrovirus. PLoS pathogens 7:e1002255.

9. Steven AC, Conway JF, Cheng N, Watts NR, Belnap DM, Harris A, Stahl SJ, Wingfield PT. 2005. Structure, assembly, and antigenicity of hepatitis B virus capsid proteins. Adv Virus Res 64:125–64.

10. Schlicksup CJ, Zlotnick A. 2020. Viral structural proteins as targets for antivirals. Curr Opin Virol 45:43–50.

11. Deres K, Schroder CH, Paessens A, Goldmann S, Hacker HJ, Weber O, Kramer T, Niewohner U, Pleiss U, Stoltefuss J, Graef E, Koletzki D, Masantschek RN, Reimann A, Jaeger R, Gross R, Beckermann B, Schlemmer KH, Haebich D, Rubsamen-Waigmann H. 2003. Inhibition of hepatitis B virus replication by drug-induced depletion of nucleocapsids. Science 299:893–6.

12. Wang XY, Wei ZM, Wu GY, Wang JH, Zhang YJ, Li J, Zhang HH, Xie XW, Wang X, Wang ZH, Wei L, Wang Y, Chen HS. 2012. In vitro inhibition of HBV replication by a novel compound, GLS4, and its efficacy against adefovir-dipivoxil-resistant HBV mutations. Antiviral therapy 17:793–803.

13. Bourne C, Lee S, Venkataiah B, Lee A, Korba B, Finn MG, Zlotnick A. 2008. Small-Molecule Effectors of Hepatitis B Virus Capsid Assembly Give Insight into Virus Life Cycle. J Virol 82:10262–10270.

14. Stray SJ, Zlotnick A. 2006. BAY 41-4109 has multiple effects on Hepatitis B virus capsid assembly. J Mol Recognit 19:542–548.

15. Schlicksup CJ, Wang JC, Francis S, Venkatakrishnan B, Turner WW, VanNieuwenhze M, Zlotnick A. 2018. Hepatitis B virus core protein allosteric modulators can distort and disrupt intact capsids. Elife 7:pii: e31473.

16. Berke JM, Dehertogh P, Vergauwen K, Mostmans W, Vandyck K, Raboisson P, Pauwels F. 2020. Antiviral Properties and Mechanism of Action Studies of the Hepatitis B Virus Capsid Assembly Modulator JNJ-56136379. Antimicrob Agents Chemother 64:pii: e02439–19.

17. Berke JM, Dehertogh P, Vergauwen K, Van Damme E, Mostmans W, Vandyck K, Pauwels F. 2017. Capsid Assembly Modulators Have a Dual Mechanism of Action in Primary Human Hepatocytes Infected with Hepatitis B Virus. Antimicrob Agents Chemother 61.

18. Wang JC, Dhason MS, Zlotnick A. 2012. Structural organization of pregenomic RNA and the carboxy-terminal domain of the capsid protein of hepatitis B virus. PLoS pathogens 8:e1002919.

19. Birnbaum F, Nassal M. 1990. Hepatitis B virus nucleocapsid assembly: primary structure requirements in the core protein. JVirol 64:3319–3330.

20. Nassal M. 1992. The arginine-rich domain of the hepatitis B virus core protein is required for pregenome encapsidation and productive viral positive-strand DNA synthesis but not for virus assembly. JVirol 66:4107–4116.

21. Lewellyn EB, Loeb DD. 2011. The arginine clusters of the carboxy-terminal domain of the core protein of hepatitis B virus make pleiotropic contributions to genome replication. Journal of Virology 85:1298–309.

22. Kann M, Schmitz A, Rabe B. 2007. Intracellular transport of hepatitis B virus. World J Gastroenterol 13:39–47.

23. Li HC, Huang EY, Su PY, Wu SY, Yang CC, Lin YS, Chang WC, Shih C. 2010. Nuclear export and import of human hepatitis B virus capsid protein and particles. PLoS pathogens 6:e1001162.

24. Yeh CT, Liaw YF, Ou JH. 1990. The arginine-rich domain of hepatitis B virus precore and core proteins contains a signal for nuclear transport. JVirol 64:6141–6147.

25. Chen C, Wang JC-Y, Zlotnick A. 2011. A kinase chaperones hepatitis B virus capsid assembly and captures capsid dynamics in vitro. PLoS pathogens 7:e1002388.

26. Chen C, Wang JC, Pierson EE, Keifer DZ, Delaleau M, Gallucci L, Cazenave C, Kann M, Jarrold MF, Zlotnick A. 2016. Importin beta Can Bind Hepatitis B Virus Core Protein and Empty Core-Like Particles and Induce Structural Changes. PLoS Pathog 12:e1005802.

27. Kann M, Sodeik B, Vlachou A, Gerlich WH, Helenius A. 1999. Phosphorylation-dependent binding of hepatitis B virus core particles to the nuclear pore complex. J Cell Biol 145:45–55.

28. Selzer L, Kant R, Wang JC, Bothner B, Zlotnick A. 2015. Hepatitis B Virus Core Protein Phosphorylation Sites Affect Capsid Stability and Transient Exposure of the C-terminal Domain. J Biol Chem 290:28584–93.

29. Zhao Q, Hu Z, Cheng J, Wu S, Luo Y, Chang J, Hu J, Guo JT. 2018. Hepatitis B virus core protein dephosphorylation occurs during pregenomic RNA encapsidation. J Virol doi:10.1128/JVI.02139-17.

30. Heger-Stevic J, Zimmermann P, Lecoq L, Bottcher B, Nassal M. 2018. Hepatitis B virus core protein phosphorylation: Identification of the SRPK1 target sites and impact of their occupancy on RNA binding and capsid structure. PLoS Pathog 14:e1007488.

31. Rabe B, Vlachou A, Pante N, Helenius A, Kann M. 2003. Nuclear import of hepatitis B virus capsids and release of the viral genome. PNAS 100:9849–9854.

32. Schmitz A, Schwarz A, Foss M, Zhou L, Rabe B, Hoellenriegel J, Stoeber M, Pante N, Kann M. 2010. Nucleoporin 153 arrests the nuclear import of hepatitis B virus capsids in the nuclear basket. PLoS Pathog 6:e1000741.

33. Rabe B, Delaleau M, Bischof A, Foss M, Sominskaya I, Pumpens P, Cazenave C, Castroviejo M, Kann M. 2009. Nuclear entry of hepatitis B virus capsids involves disintegration to protein dimers followed by nuclear reassociation to capsids. PLoS Pathog 5:e1000563.

34. Kann M, Thomssen R, Kochel HG, Gerlich WH. 1993. Characterization of the endogenous protein kinase activity of the hepatitis B virus. Arch Virol Suppl 8:53–62.

35. Osseman Q, Kann M. 2017. Intracytoplasmic Transport of Hepatitis B Virus Capsids. Methods Mol Biol 1540:37–51.

36. Cingolani G, Petosa C, Weis K, Muller CW. 1999. Structure of importin-beta bound to the IBB domain of importin-alpha. Nature 399:221–9.

37. Hilmer JK, Zlotnick A, Bothner B. 2008. Conformational Equilibria and Rates of Localized Motion within Hepatitis B Virus Capsids. J Mol Biol 375:581–594.

38. Porterfield JZ, Dhason MS, Loeb DD, Nassal M, Stray SJ, Zlotnick A. 2010. Full-Length Hepatitis B Virus Core Protein Packages Viral and Heterologous RNA with Similarly High Levels of Cooperativity. Journal of Virology 84:7174–7184.

39. Chua PK, Tang FM, Huang JY, Suen CS, Shih C. 2010. Testing the balanced electrostatic interaction hypothesis of hepatitis B virus DNA synthesis by using an in vivo charge rebalance approach. Journal of Virology 84:2340–51.

40. Schlicksup CJ, Laughlin P, Dunkelbarger S, Wang JC, Zlotnick A. 2020. Local Stabilization of Subunit-Subunit Contacts Causes Global Destabilization of Hepatitis B Virus Capsids. ACS Chem Biol 15:1708–1717.

41. Rayleigh L. 1882. XX. On the equilibrium of liquid conducting masses charged with electricity. Philos Mag 14:184–186.

42. Lutomski CA, Gordon SM, Remaley AT, Jarrold MF. 2018. Resolution of Lipoprotein Subclasses by Charge Detection Mass Spectrometry. Anal Chem 90:6353–6356.

43. Cui X, Ludgate L, Ning X, Hu J. 2013. Maturation-associated destabilization of hepatitis B virus nucleocapsid. J Virol 87:11494–503.

44. Dhason MS, Wang JC, Hagan MF, Zlotnick A. 2012. Differential assembly of Hepatitis B Virus core protein on single- and double-stranded nucleic acid suggest the dsDNA-filled core is spring-loaded. Virology 430:20–9.

45. Stray SJ, Bourne CR, Punna S, Lewis WG, Finn MG, Zlotnick A. 2005. A heteroaryldihydropyrimidine activates and can misdirect hepatitis B virus capsid assembly. Proc Natl Acad Sci U S A 102:8138–43.

46. Venkatakrishnan B, Katen SP, Francis S, Chirapu S, Finn MG, Zlotnick A. 2016. Hepatitis B Virus Capsids Have Diverse Structural Responses to Small-Molecule Ligands Bound to the Heteroaryldihydropyrimidine Pocket. J Virol 90:3994–4004.

47. Rudnick J, Bruinsma R. 2005. Icosahedral packing of RNA viral genomes. Phys Rev Lett 94:038101.

48. Nair S, Zlotnick A. 2018. Asymmetric Modification of Hepatitis B Virus (HBV) Genomes by an Endogenous Cytidine Deaminase inside HBV Cores Informs a Model of Reverse Transcription. J Virol 92:pii: e02190–17.

49. Dykeman EC, Grayson NE, Toropova K, Ranson NA, Stockley PG, Twarock R. 2011. Simple rules for efficient assembly predict the layout of a packaged viral RNA. Journal of molecular biology 408:399–407.

50. Yau ST, Thomas BR, Galkin O, Gliko O, Vekilov PG. 2001. Molecular mechanisms of microheterogeneity-induced defect formation in ferritin crystallization. Proteins 43:343–52.

51. Katen SP, Tan Z, Chirapu SR, Finn MG, Zlotnick A. 2013. Assembly-directed antivirals differentially bind quasiequivalent pockets to modify hepatitis B virus capsid tertiary and quaternary structure. Structure 21:1406–16.

52. Seeger C, Mason WS. 2015. Molecular biology of hepatitis B virus infection. Virology 479–480:672–86.

53. Sun L, Wu J, Du F, Chen X, Chen ZJ. 2013. Cyclic GMP-AMP synthase is a cytosolic DNA sensor that activates the type I interferon pathway. Science 339:786–91.

54. Guo F, Zhao Q, Sheraz M, Cheng J, Qi Y, Su Q, Cuconati A, Wei L, Du Y, Li W, Chang J, Guo JT. 2017. HBV core protein allosteric modulators differentially alter cccDNA biosynthesis from de novo infection and intracellular amplification pathways. PLoS Pathog 13:e1006658.

55. Chopra A, Willmore WG, Biggar KK. 2019. Protein quantification and visualization via ultraviolet-dependent labeling with 2,2,2-trichloroethanol. Sci Rep 9:13923.

56. Schindelin J, Arganda-Carreras I, Frise E, Kaynig V, Longair M, Pietzsch T, Preibisch S, Rueden C, Saalfeld S, Schmid B, Tinevez JY, White DJ, Hartenstein V, Eliceiri K, Tomancak P, Cardona A. 2012. Fiji: an open-source platform for biological-image analysis. Nat Methods 9:676–82.

57. Burns EE, Keith BK, Refai MY, Bothner B, Dyer WE. 2017. Proteomic and biochemical assays of glutathione-related proteins in susceptible and multiple herbicide resistant Avena fatua L. Pestic Biochem Physiol 140:69–78.

58. Ledbetter RN, Garcia Costas AM, Lubner CE, Mulder DW, Tokmina-Lukaszewska M, Artz JH, Patterson A, Magnuson TS, Jay ZJ, Duan HD, Miller J, Plunkett MH, Hoben JP, Barney BM, Carlson RP, Miller AF, Bothner B, King PW, Peters JW, Seefeldt LC. 2017. The Electron Bifurcating FixABCX Protein Complex from Azotobacter vinelandii: Generation of Low-Potential Reducing Equivalents for Nitrogenase Catalysis. Biochemistry 56:4177–4190.

59. Vaudel M, Burkhart JM, Zahedi RP, Oveland E, Berven FS, Sickmann A, Martens L, Barsnes H. 2015. PeptideShaker enables reanalysis of MS-derived proteomics data sets. Nat Biotechnol 33:22–4.

60. Barsnes H, Vaudel M. 2018. SearchGUI: A Highly Adaptable Common Interface for Proteomics Search and de Novo Engines. J Proteome Res 17:2552–2555.

61. Kremer JR, Mastronarde DN, McIntosh JR. 1996. Computer visualization of three-dimensional image data using IMOD. J Struct Biol 116:71–6.

62. Frangakis AS, Hegerl R. 2001. Noise reduction in electron tomographic reconstructions using nonlinear anisotropic diffusion. J Struct Biol 135:239–50.

63. Contino NC, Jarrold MF. 2013. Charge detection mass spectrometry for single ions with a limit of detection of 30 charges. Int J Mass Spect 345–347:153–159.

64. Draper BE, Anthony SN, Jarrold MF. 2018. The FUNPET-a New Hybrid Ion Funnel-Ion Carpet Atmospheric Pressure Interface for the Simultaneous Transmission of a Broad Mass Range. J Am Soc Mass Spectrom 29:2160–2172.

65. Draper BE, Jarrold MF. 2019. Real-Time Analysis and Signal Optimization for Charge Detection Mass Spectrometry. Journal of The American Society for Mass Spectrometry 30:898–904.

66. de la Mora JF. 2000. Electrospray ionization of large multiply charged species proceeds via Dole’s charge residue mechanism. Anal Chim Acta 406:93–104.

67. Konermann L, Ahadi E, Rodriguez AD, Vahidi S. 2013. Unraveling the mechanism of electrospray ionization. Anal Chem 85:2–9.

68. Keifer D, Motwani T, Teschke CM, Jarrold MF. 2016. Acquiring structural information on virus particles via charge detection mass spectrometry. J Am Soc Mass Spectrom 27:1028–1036.

